# Spatio-temporal dynamics of stress-induced network reconfigurations reflect negative affectivity

**DOI:** 10.1101/2021.07.16.452622

**Authors:** Anne Kühnel, Michael Czisch, Philipp G. Sämann, BeCOME Study Team, Elisabeth B. Binder, Nils B. Kroemer

**Author notes:** Corresponding authors: A. Kühnel, E. B. Binder. contributed equally (Elisabeth B. Binder, Nils B. Kroemer).

## Abstract

**Background:** Chronic stress is an important risk factor in the etiology of mood and anxiety disorders, but exact pathomechanisms remain to be understood. Mapping individual differences of acute stress-induced neurophysiological changes, especially on the level of neural activation and functional connectivity (FC), could provide important insights in how variation in the individual stress response is linked to disease risk.

**Methods:** Using an established psycho-social stress task flanked by two resting-state scans, we measured subjective, physiological, and brain responses to acute stress and recovery in 217 unmedicated participants with and without mood and anxiety disorders. To estimate block-wise changes in stress-induced brain activation and FC, we used hierarchical mixed-effects models based on denoised timeseries within a predefined stress network. We predicted inter- and intra-individual differences in stress phases (anticipation vs. acute stress vs. recovery) and transdiagnostic dimensions of stress reactivity using elastic net and support vector machines.

**Results:** We identified four subnetworks showing distinct changes in FC over time. Subnetwork trajectories predicted the stress phase (accuracy: 71%, *p*_perm_<.001) and increases in pulse rate (*R^2^*=.10, *p*_perm_<.001). Critically, individual spatio-temporal trajectories of changes across networks also predicted negative affectivity (Δ*R^2^*=.08, *p*_perm_=.009), but not the presence or absence of a mood and anxiety disorder.

**Conclusions:** Spatio-temporal dynamics of brain network reconfiguration induced by stress reflect individual differences in the psychopathology dimension negative affectivity. These results support the idea that vulnerability for mood and anxiety disorders can be conceptualized best at the level of network dynamics, which may pave the way for improved prediction of individual risk.

## 1. Introduction

Stressful situations occur frequently in everyday life and an adaptive response to stress is critical for mental health (1). Congruently, maladaptive stress responses such as prolonged anxiety, extensive rumination, and negative coping strategies are common symptoms of many mental disorders including mood and anxiety disorders (2–5). These maladaptive responses to stress are mirrored on a biological level, where dysregulation of the endocrine (6–9) as well as autonomous stress response (10) have been described across mood and anxiety disorders.

The stress response can be divided into three phases: anticipation (11, 12), the acute (13, 14) stress response and recovery (15–17). All phases have been shown to be affected on different levels in relation with risk to mood and anxiety disorders (13, 14). An increased endocrine responses in anticipation of stress (18–20), a blunted acute response (9), or prolonged recovery after stress (21) have been associated with depression. Depression-related personality characteristics such as negative affectivity or trait anxiety (22, 23) with shared genetic signatures (24) also affect endocrine stress reactivity across all phases (25–27). Likewise, mood and anxiety disorders are characterized by specific maladaptive cognitions related to stress (28, 29). For instance, negative coping styles, such as excessive rumination, often seen in depressed patients, are associated with slower stress recovery (30–33), whereas distraction is associated with faster recovery (34, 35). Resilient coping styles, such as social support (36) or cognitive reappraisal (37), showed faster stress recovery (38) and reduced anticipatory stress (39). Taken together, specific psychological factors of mood and anxiety disorders may alter stress reactivity at different phases of the stress response.

On the neural level, stress responses are characterized by dynamic shifts in the salience (SN), default mode (DMN), and fronto-parietal (FPN) networks (40, 41). Consequently, changes in FC between key nodes of the stress network have been reported (42, 43) up to 40min after stress. Within this network, dysregulation has been consistently reported across mood and anxiety disorders (44), suggesting that brain networks implicated in acute stress reactivity are also affected in disorders showing maladaptive stress responses. Comparably, previous work in healthy participants or adolescents with mental disorders has shown that trait anxiety is associated with altered stress-induced activation in key regions of the stress network (45, 46). However, most studies focus on average stress-induced brain responses during the task. Hence, little is known about dynamic changes within the stress network across the stress cycle although emerging evidence has highlighted the importance dynamic reconfigurations of brain networks in mental disorders (47, 48). Likewise, task-induced changes in FC have been shown to improve correspondence with phenotypic differences compared to resting FC and have been proposed as promising target to unravel alterations in mental disorders (49, 50). Therefore, identifying individual signatures of stress reactivity that may map to risk for psychopathology could help pinpoint potential intervention targets (i.e., for non-invasive brain stimulation techniques (51) and improve means to study network perturbations in clinical trials.

Here, we used a hierarchical model of stress-induced changes in brain responses and FC to characterize trajectories of network reconfigurations across the stress cycle. Using individual FC signatures of stress adaptation, we identified dynamic FC changes that differentiate between stress states and predict interindividual differences in negative affectivity, providing a link between acute psycho-social stress reactivity and psychopathology.

## 2. Methods

### 2.1 Participants

The sample was recruited as part of the Biological Classification of Mental Disorders study at the Max Planck Institute of Psychiatry (ClinicalTrials.gov: NCT03984084,(52)). The study characterizes participants with a broad spectrum of mood and anxiety disorders including common comorbidities as well as unaffected individuals. Here, we included 217 participants (140 women, M_age_= 35.1 years ± 12.1, Supplementary Table S1). All participants underwent a computer-based, standardized diagnostic interview (CIDI (53)). Diagnoses were derived by a DSM-IV-based algorithm and n=129 (54%) fulfilled the criteria for ≥1 mood or anxiety disorder (ICD-10 code F3-F4, excluding specific phobias) within the last 12 months (Supplementary Table S2). None of the participants reported any present medication for their psychiatric symptoms. To maximize the sample size, we excluded participants with missing or low-quality data for each analysis separately leading to sample sizes between 177 and 217 (Supplementary Table S4).

### 2.2 Experimental procedure

The imaging stress task (Supplementary Figure S1) was included in the second functional magnetic resonance imaging (fMRI) session (52), so none of the participants were fMRI naïve (54). Upon arrival, the first saliva sample was taken (T0) for cortisol assessment followed by a second sample (T1) approximately 20 min later, after placement of an intravenous catheter for additional blood sampling in 73 (33%) participants, and before entering the scanner. After an emotional face-matching task (∼12 min), a baseline resting-state measurement, and immediately before the stress paradigm, participants rated their current affective state using *Befindensskalierung nach Kategorien und Eigenschaftsworten* (BSKE, (55) Supplementary Information). The psycho-social stress paradigm was adapted from the Montreal imaging stress task (56), where stress is induced by performing arithmetic tasks under time pressure and with negative feedback (57, 58). The task lasted ∼25 min and included a *PreStress* phase without negative feedback or time pressure, followed by a *Stress* phase with psycho-social stress-induction, and a *PostStress* phase (analogous to *PreStress*). Each phase contained 5 task blocks (50s) interleaved with rest (40s) blocks. We measured autonomous activity throughout using photoplethysmography (Supplementary Information). After completion of the task, affective state was assessed and saliva samples were taken (T6). A 30-min rest period lying outside the scanner was followed by a concluding resting-state scan, and assessments of subjective affect and saliva cortisol (T8). In participants with additional blood sampling, samples were taken in the scanner before the task (T3), during, and after the task in approximately 15-min intervals (T4-T8). At the end of the session, participants were debriefed.

### 2.3 Questionnaires

To measure state- and trait-like depressive symptoms and negative affect (23), we included the *Becks Depression Inventory-II* (BDI, (59)) and the trait subscale of the *State-Trait Anxiety Inventory* (TAI, (60)). To measure maladaptive and adaptive psychological stress reactivity, we included the *Intolerance of uncertainty scale* (IoU, (61)), a stress coping scale (*Stressverarbeitungsfragebogen,* SVF,(62)), and a resilience scale (RS-11,(63)).

### 2.4 fMRI data acquisition and preprocessing

Briefly, MRI data were acquired on a GE 3T scanner (Discovery MR750, GE, Milwaukee, U.S.A.). The functional data were 755 T2*-weighted echo-planar images (EPI) for the stress task and 155 EPIs for each of the resting-state scans. All fMRI data preprocessing was performed in Matlab 2018a (The Mathworks Inc., Natick, MA, USA) and SPM12 (v12; Wellcome Department of Imaging Neuroscience, London, UK). Data was slice-time corrected, realigned, normalized to the MNI-template using DARTEL (64), and spatially smoothed with a 6×6×6mm^3^ full-width at half maximum kernel (Supplementary Information).

### 2.5 Data analysis

#### 2.5.1 Questionnaire data: Non-negative matrix factorization

To extract interpretable dimensions capturing maladaptive stress reactivity from the questionnaires, we used non-negative matrix factorization (NNMF, (65)). In contrast to other dimension reduction methods, NNMF captures additive latent variables that are intuitively interpretable since all weights are positive. We included all items from the questionnaires after rescaling them between 0 and 1. We estimated NNMF (*nnmf*, MATLAB 2020b) with 150 iterations and 50,000 replicates to ensure stability. To determine the optimal number of dimensions, we used the elbow method for explained variance (66).

#### 2.5.2 Stress response to the psycho-social stress task

The endocrine stress response was estimated as the change in cortisol concentration (ΔCort) between T1 and T6. Since we took blood samples in a subset of participants and a cortisol response to the procedure may confound the response to the stress task (58), we included a nuisance regressor (dummy coded) classifying participants with a response > 2.5 nmol/l (0.91 ng/ml, (58, 67)) at T1 (20 min after arrival) compared to baseline (T0) as pre-task cortisol responders in all analyses.

The autonomous stress response was estimated as change in average pulse rate during the arithmetic blocks in the *Stress* phase compared to *PreStress* or *PostStress*, respectively (58). Subjective stress effects were estimated as the change in positive and negative affect (sum scores across 15 BSKE items) after the task (57, 58) (Supplementary Information).

#### 2.5.3 fMRI data

To compare stress-induced changes in brain response with previous studies, we assessed associations between stress-induced changes in activation and dimensions of stress susceptibility using previously reported first-level contrasts including one task regressor for each task phase (*PreStress*, *Stress*, and *PostStress*) phase (58) (Supplementary Information). At the group-level, we used voxel-wise multiple regressions. To capture stress-induced changes independent of directionality, we calculated representational similarity (58, 68) (Supplementary Information) between average contrasts of *PreStress*, *Stress*, and *PostStress* phases in 268 atlas ROIs (69). Individual similarity was calculated as Fisher’s z-transformed Pearson correlations. All analyses regarding fMRI data and psychometric variables (whole-brain regressions and elastic net predictions) included age, sex, pre-task cortisol, and average log-transformed framewise displacement as covariates (Supplementary Information).

To model dynamic changes in FC across the task, we extracted average timeseries from the preprocessed but unsmoothed stress task, the preceding and the following resting state in 21 ROIs based on previously reported activations (Fig. 1B, (40,41,45,70)). The regions included the left and right amygdala, hypothalamus, caudate, putamen, anterior, medial, and posterior hippocampus, anterior and posterior insula and one region for the posterior cingulate, dorsal anterior cingulate, and ventromedial prefrontal cortex. Regions were defined using a FC-based atlas (69), only the hypothalamus was defined based on an the Harvard-Oxford atlas as the resolution of the Shen atlas was too coarse. Timeseries were detrended (linear), despiked (winsorized at ±4SDs), and residualized with the same covariates as in previous work (57) including the 6 movement parameters, their derivative, and 5 components extracted from white matter and cerebro-spinal fluid, respectively ((71), Supplementary Information). To estimate changes relative to the resting-state baseline before the task, we concatenated all data by matching their grayscale values (Supplementary Information).

**Figure 1:**
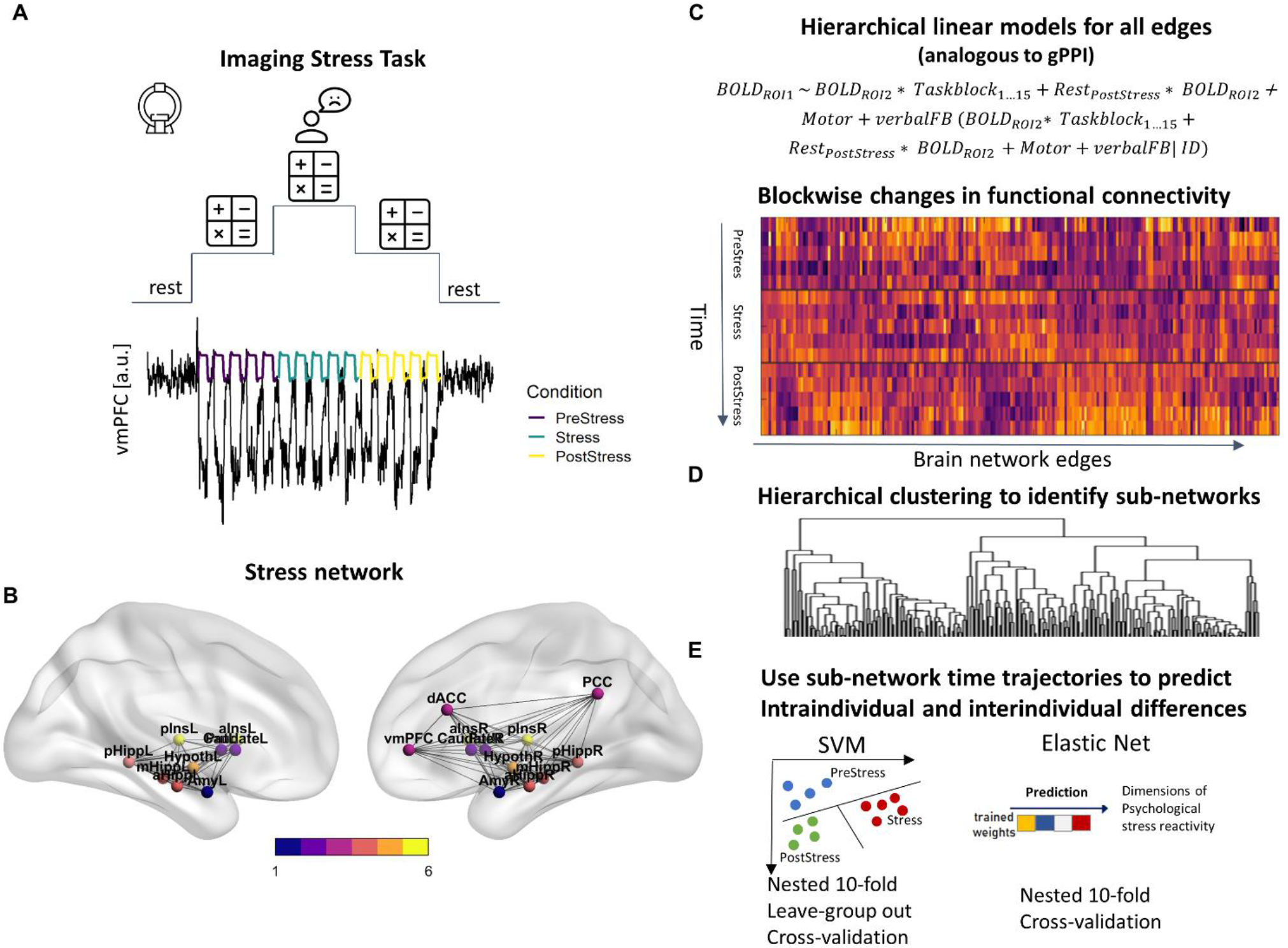
Schematic overview of task design and analyses. A) The psycho-social stress task consists of 15 blocks (50s each) of arithmetic problems interleaved with rest blocks (fixation cross, 40s each). The first five blocks are without aversive feedback (*PreStress*), followed by five blocks with negative feedback and time constraints (*Stress),* and another five blocks without aversive feedback (*PostStress*). Illustratively, we depict the average time series of the vmPFC after denoising across all measurements, which tracks the structure of the paradigm. B) Stress-induced changes in activation and functional connectivity (FC) from block to block are characterized for all regions and edges within a predefined stress network. C) Changes in activation and FC for each block are estimated using a hierarchical extension of generalized psychophysiological interactions (gPPI) estimated with one hierarchical linear model for each edge of the network, leading to group-level estimates of task-induced FC change for each block and 210 edges. D) For further predictive analyses, edges with similar changes over time are clustered into four subnetworks using hierarchical clustering. E) Lastly, we use individual-level profiles of the four subnetworks FC changes (average across all edges per subnetwork) to predict either the task phase of unseen blocks (four features per block) or interindividual differences in adaptive vs. maladaptive stress reactivity, vmPFC = ventromedial prefrontal cortex, dACC = dorsal anterior cingulate cortex, Put = putamen, PCC = posterior cingulate cortex, pIns = posterior insula, aIns = anterior insula, pHipp = posterior hippocampus, mHipp = medial hippocampus, aHipp = anterior hippocampus, Amy = amygdala, SVM = support vector machine

We then used hierarchical linear models (LME, (72, 73)) analogous to hierarchical generalized psychophysiological interactions (74) to estimate block-wise changes in activation in all 21 ROIs (Fig. 1B) and FC for all (21*20)/2 edges between nodes. We estimated one model for each edge with all predictors as random effects, deriving group-level and regularized individual-level estimates simultaneously (75–77). Each model included the timeseries of one region (ROI_1_) as a dependent variable and the timeseries of the other region (ROI_2_) as independent variable together with one regressor for each of the 15 task blocks (convolved with hemodynamic response function) and the interaction of each task-block regressor with the predictive timeseries. Additionally, we included an interaction term for the post task resting-state to account for lasting stress-induced changes in FC. For interaction terms, the predictors were mean-centered.

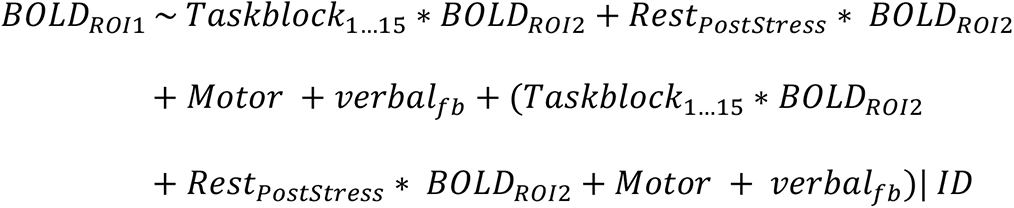

To reduce dimensionality, we defined subnetworks of nodes showing similar FC changes over time (i.e., blocks) using hierarchical clustering (*eclust,* (78)) with z-standardization and Pearson correlation as distance measure. The number of clusters was determined by evaluating the decrease in total within sum-of-squares (*wss*) with the elbow method leading to 4 distinct clusters (Supplementary Figure S5).

To evaluate the predictive performance of stress-induced FC changes within the subnetworks, we used machine learning algorithms to predict intra- and inter-individual differences in stress susceptibility. First, we predicted the task phase (*PreStress*, *Stress*, or *PostStress*) of unseen blocks based on average FC-changes in the 4 subnetworks using support-vector machine (SVM) classifiers with a radial basis function (one vs. one, *SVC,* scikitlearn (79), Python 3.7.0) with nested 10-fold cross-validation. We used a leave-subject-out approach so that all data from 10% of the participants was in a held-out fold. Second, we used the same approach to predict z-scored pulse-rate changes for each block using support-vector regression (SVR). To test whether the prediction provided information in addition to differences between stress phases (i.e., higher pulse rate during stress), we estimated LMEs including the observed pulse-rate change of each block, the stress phase and their interaction as predictors (random effects by participant, (80)). Last, we predicted interindividual differences in dimensions of stress reactivity derived from NNMF using activation (12 ROIs) and connectivity (4 subnetworks) trajectories across the task blocks. Since the models included between 68 (connectivity) and 180 (activation) features, we used elastic net (*lasso*, preset alpha = .5) with nested 10-fold cross-validation. Elastic net performs well if features are correlated and their number is moderately high compared to the number of observations (81). To account for confounding variables, we included them in the baseline prediction models and evaluated the incremental variance explained by fMRI features. Notably, average log-transformed framewise displacement was not associated with diagnosis status or psychometric dimensions of stress reactivity (*r*s < .12, *p*s > .11). Statistical significance was determined using permutation tests (1,000 iterations), where the outcome was shuffled together with the confounders to keep their correlation structure.

#### 2.5.4 Statistical threshold and software

Statistical analyses were performed in R v3.5.1. (82). For whole-brain fMRI analyses, the voxel threshold was set at *p*<.001 (uncorrected). Clusters were considered significant with an FWE cluster-corrected *p*-value threshold of *p_cluster.FWE_*<.05. Additional LME models were estimated in R using lmerTest (83).

## 3. Results

### Average task-induced stress responses do not differ in mood and anxiety disorders

The task induced stress across multiple levels: positive affect decreased (*b*=-2.35, *p*<.001), while negative affect (*b*=7.6, *p*<.001, Fig. 2A) increased after the task. Likewise, pulse rate (*b*=6.5, *p*<.001, Fig. 2B) increased during stress as well as salivary cortisol (*b*=.42, *p*=.007, Fig. 2C). On the neural level, stress led to significant deactivation in the DMN (PCC and angular gyrus), insula, dorsomedial prefrontal cortex as well as activation in the visual and parietal cortex (see Fig. 2E).

**Figure 2:**
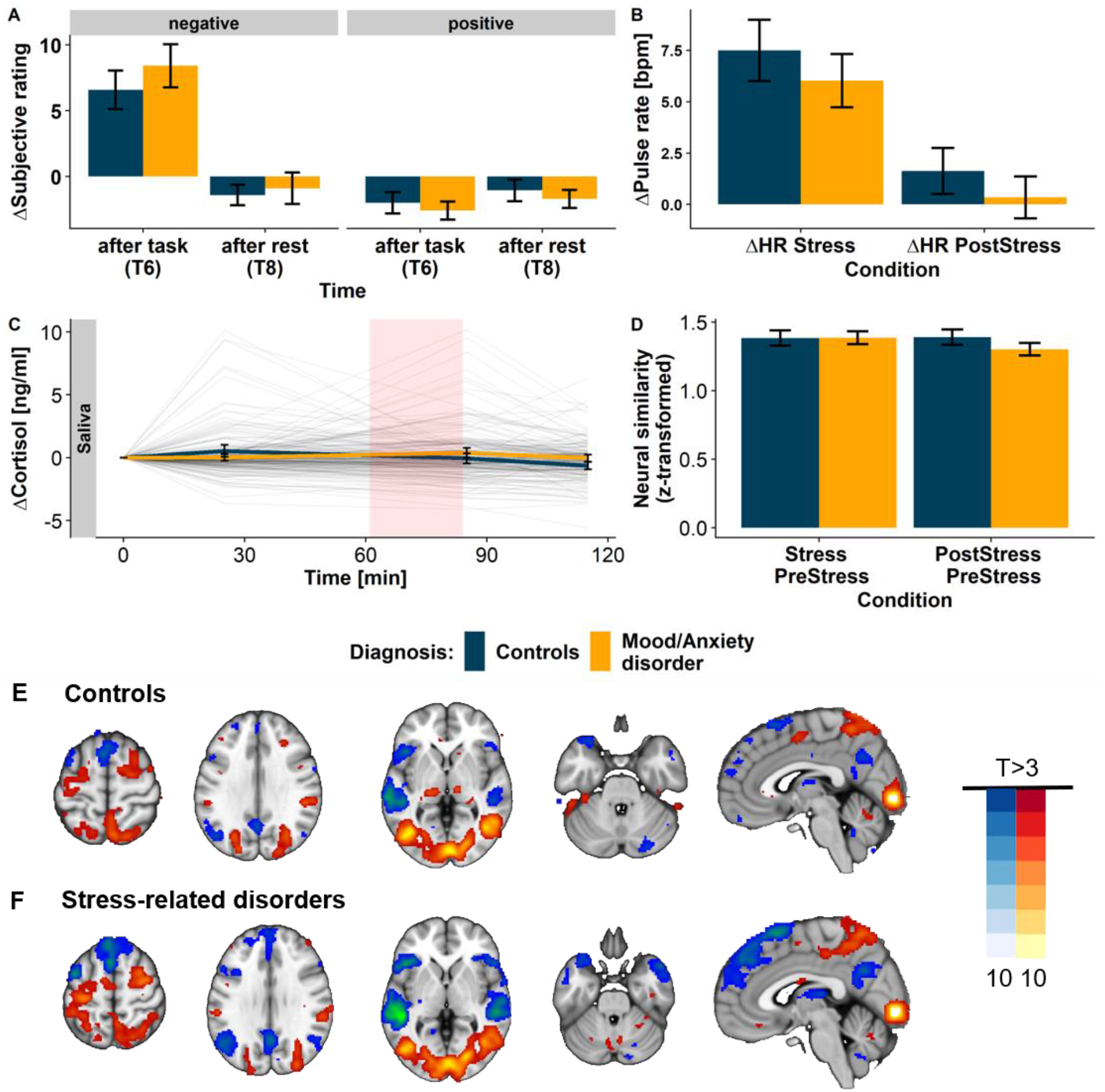
The psycho-social stress task leads to multi-modal stress responses that do not differ in participants with mood and anxiety disorders. A) Pulse rate increases during the *Stress* phase and recovers in the *PostStress* phase similarly in both groups. B) Negative affect increases (T6: *b*=-7.6, *p*<.001) and recovers after stress (T8: *b* = −1.1, *p* = .006), while positive affect decreases (T6: *b*=2.35, *p*<.001, Fig. 2A) and does not recover back to baseline levels (T8: *b* = −1.4, *p* < .001) in both groups (Supplementary Table S3). C) The task leads to an increase in salivary cortisol compared to baseline (T0). Thin lines depict individual cortisol trajectories, thick lines show group averages. The shaded area shows the timing of the stress task. D) Neural similarity (58) assessing regional and directional unspecific neural changes comparing *PostStress* but not *Stress* (*b*=0.0, *p*=.80) to the *PreStress* baseline differed between groups. E) Stress-induced changes in neural activation do not differ between groups. All models include age, sex, and pre-task cortisol response as covariates and response variables are residualized accordingly. Neural similarity models additionally included average framewise displacement. Error bars depict 95% confidence intervals.

In contrast to previous reports, average stress reactivity did not differ between participants with and without mood and anxiety disorders on the autonomous, endocrine, or subjective level (Fig. 2, Supplementary Table S3). Likewise, there were no significant differences in brain response on average whole-brain maps (Fig. 2E), although neural similarity during stress recovery was significantly lower in participants with a mood or anxiety disorder (*b*=-0.05, *p*=.005, Fig. 2B), indicating slower recovery. The lack of significant alterations of stress reactivity may be due to the heterogeneous phenotype of the patient group, or because individual stress-induced changes are only insufficiently reflected in average maps that lack dynamic information about network-level reconfiguration.

### Dynamic connectivity changes predict stress state and changes in pulse rate

To assess stress-induced changes throughout the stress cycle, we used concatenated data from the psycho-social stress task and two flanking resting-state scans (see Fig. S1). To derive stress-induced changes at a single-block resolution, we used mixed-effects models of fMRI timeseries. By fitting a hierarchical model, such estimates recover individual deviations from group averages more robustly (75–77). While stress-induced changes in brain responses were similar across task blocks (Supplementary Figure S3), FC changes were qualitatively and quantitatively discernable across stress phases (Fig. 3B). To reduce dimensions for individual predictions, we identified subnetworks of edges with a comparable stress response using hierarchical clustering. According to the elbow criterion (*wss*), we identified four clusters of subnetworks showing distinct stress-induced changes (Figure 3, Supplementary Figure S4-6). Two clusters showed a pronounced FC change in response to stress onset that was followed by gradual recovery towards the baseline state: the blue cluster, primarily reflecting cross-clique connections (i.e., between canonical networks), and the yellow cluster, primarily reflecting cortico-limbic connections. In contrast, the green cluster, primarily including DMN edges, showed decreasing FC and the purple cluster, primarily including edges from the hypothalamus, showed increasing FC throughout the complete task, suggesting that its FC does not recover to the *PreStress* state.

**Figure 3:**
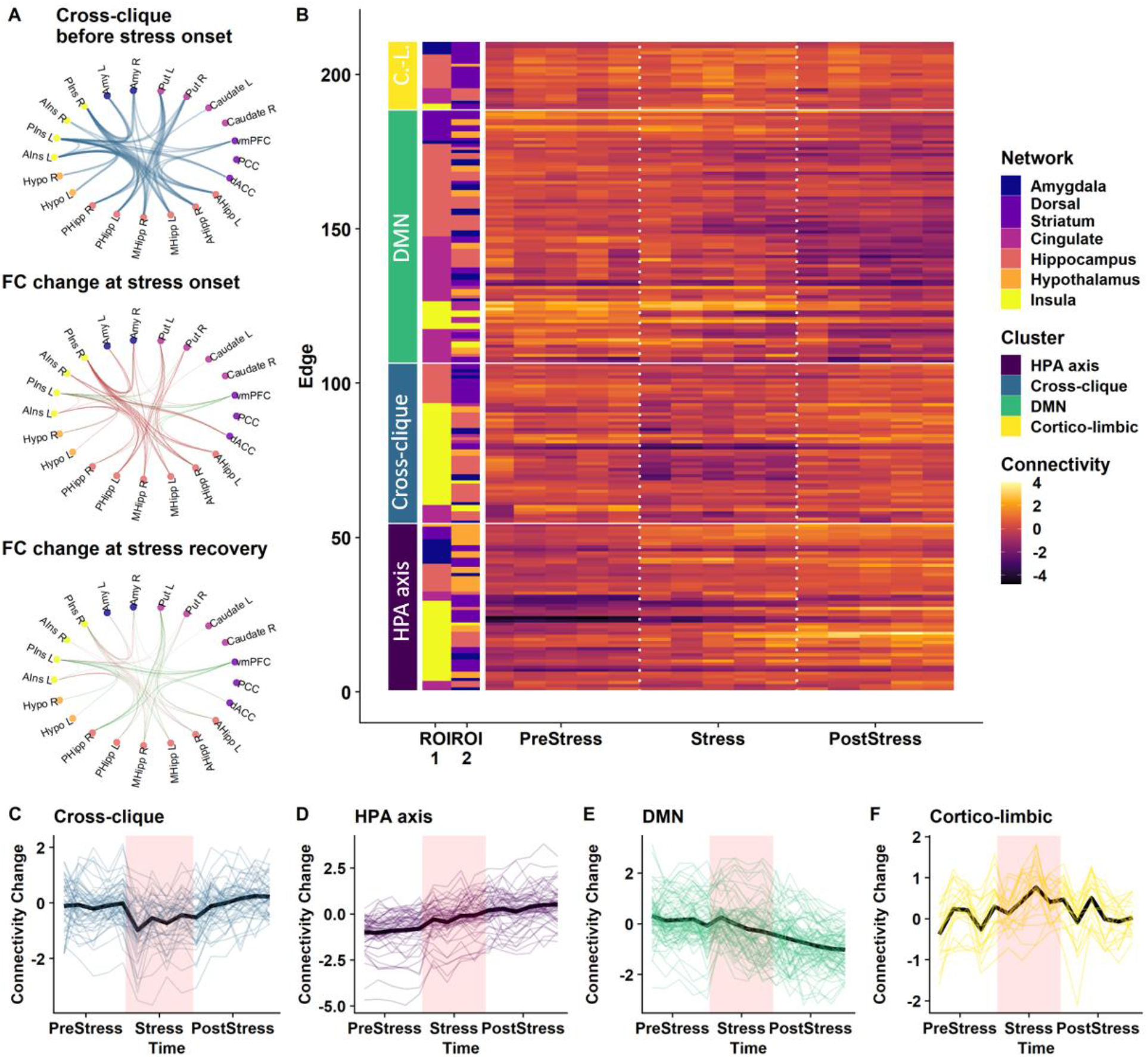
Psycho-social stress leads to characteristic spatio-temporal patterns of functional connectivity (FC) changes. A: A cluster reflecting cross-clique connections shows a decrease in FC in response to stress and slowly recovers afterwards. The first circle plot shows the change in FC strength in the first block compared to rest (standardized and rescaled for visualization) for all edges. The second and third plot show the change in FC compared to the beginning of the task at stress onset (first stress block) and at the end of stress recovery (last *PostStress* block). Red lines indicate decreases in FC and green lines increases, line thickness shows the strength of change. The circle plots for the other 3 networks are shown in the Figure S6. B: FC change (z-standardized) in edges of the stress network ordered according to the subnetworks identified by hierarchical clustering. C-F) Trajectories of block-wise FC changes (z-standardized) for all four subnetworks (thin lines depict individual edges, thick lines the average across all edges of the subnetwork). vmPFC = ventromedial prefrontal cortex, dACC = dorsal anterior cingulate cortex, Put = putamen, PCC = posterior cingulate cortex, pIns = posterior insula, aIns = anterior insula, pHipp = posterior hippocampus, mHipp = medial hippocampus, aHipp = anterior hippocampus, Amy = amygdala, DMN = default mode network, HPA axis = Hypothalamus-pituitary-adrenal axis

Next, to verify that these spatio-temporal profiles indeed reflect the experimentally induced stress phases, we predicted the phase of unseen blocks based on individual-level estimates within the four subnetworks using SVM. Stress-induced changes in FC predicted stress phases with high accuracy (71% vs. 33% chance; *p_perm_*<.001, individual accuracy *M*=71% ±14%, Figure 4A). However, predictions solely based on changes in activation barely exceeded chance levels (40%, Figure 4B). The same FC features predicted relative changes in pulse rate of each block within participants using SVR (*r*=.31, *R*^2^=.10, *p_perm_*<.001, Figure 4C). Critically, successful prediction of pulse rate was not only driven by changes between task phases (e.g., higher pulse rate during stress), but also recovered differences in pulse rate of blocks within the same stress phase (*p*s between .02 and <.001, Fig. 4D, Supplementary Information). Stress-induced increases in pulse rate (*Stress – PreStress*) derived from predicted changes in pulse rate for each block corresponded with observed stress-induced effects (*p*s≤.002, Fig. 4E-F). Decreasing or further increasing the number of subnetworks derived from the hierarchical clustering did not improve the predictive performance (Supplementary Figure S7). Changes in head movement during stress alone could not explain the successful prediction of stress phases, since a prediction based on average framewise-wise displacement and average differences in consequent images (DVARS) of each task block performed significantly worse (43%, Supplementary Information, Figure S11) To summarize, spatio-temporal profiles of stress-induced responses within the four subnetworks track the current stress phase and physiological adaptation better than chance or changes in brain response.

**Figure 4:**
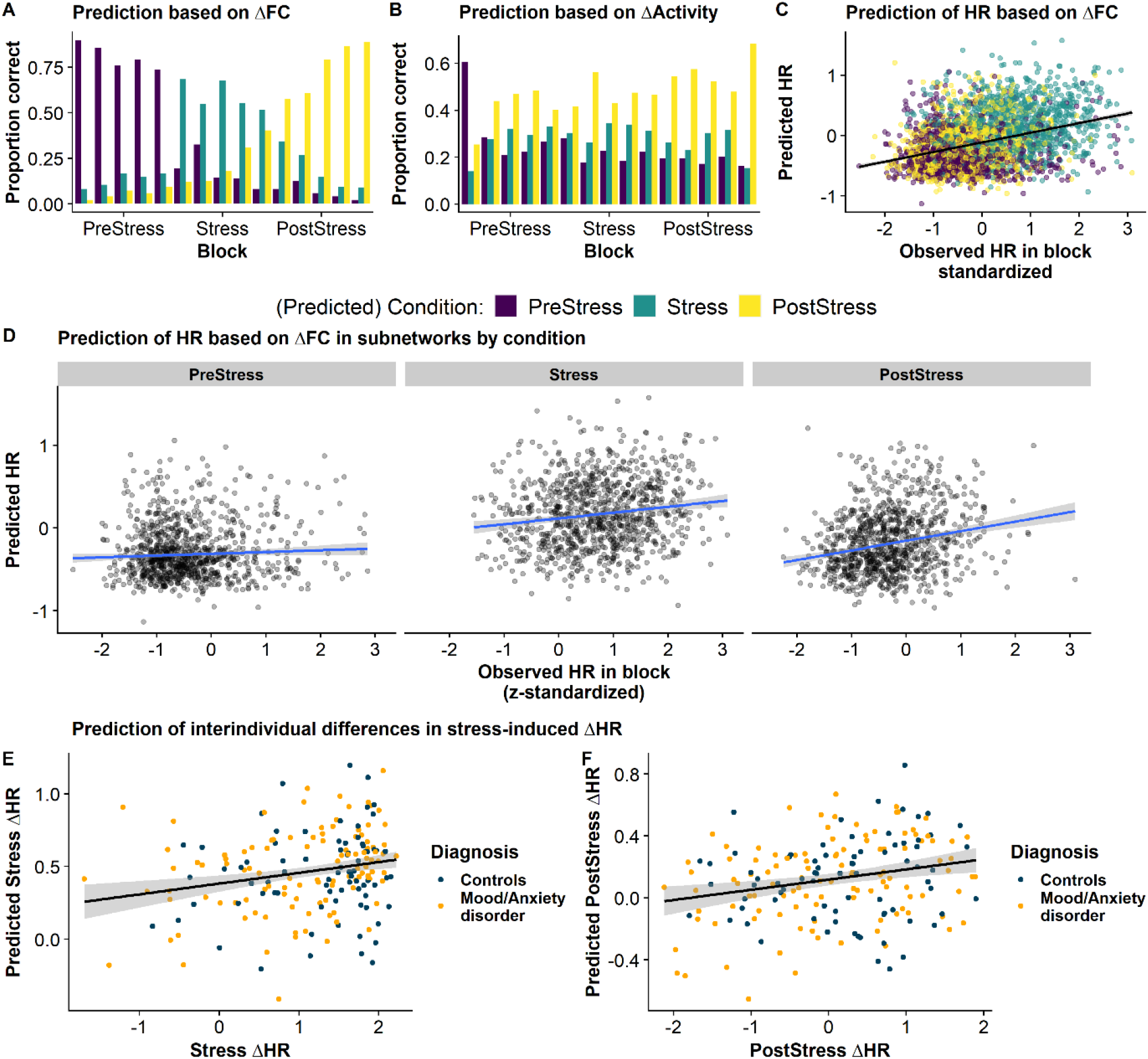
Block-wise changes in functional connectivity (FC) in the four stress subnetworks predict the stress state and individual changes in pulse rate in unseen blocks. A: Block-wise changes in FC predict the current stress phase above chance (71%, *p*_perm_<.001). Predictions are best for the *PreStress* condition and the initial stress blocks. In contrast, the transition from *Stress* to *PostStress* is harder to differentiate, indicating a gradual recovery into discernable states of recovery. B: Predictions based solely on changes in brain responses do not exceed chance levels (40%). C: Changes in FC predict changes in pulse rate within participants (*R*^2^=.10, *p*<.001). To account for baseline differences, the pulse is z-standardized within each participant. D: Successful prediction of changes in pulse rate does not only recover differences between stress and non-stress conditions, but also predicts the magnitude of pulse-rate changes within stress recovery and acute stress phases. E-F: Comparing inter-individual differences in stress-induced changes (e.g., *Stress-PreStress* (*r*=.22, *p*=.002) or *PostStress-PreStress* (*r*=.25, *p*<.001)), derived from the observed and the predicted pulse rate changes of each block, showed significant correlations, indicating that inter-individual differences in the stress-induced pulse rate response can also be recovered.

### Dynamic connectivity changes predict negative affectivity as a transdiagnostic dimension reflecting maladaptive stress reactivity

To map differences in dynamic FC to psychological constructs related to stress adaptation, we derived questionnaire-based dimensions reflecting psychological responses to stress (i.e., resilience and susceptibility) using NNMF. To this end, we included single-item responses assessing state and trait factors including depressive symptoms (BDI), trait-like negative affect (TAI) as well as stress coping (SVF), intolerance of uncertainty (IoU) and resilience (Resilience). The most parsimonious solution revealed five interpretable dimensions (Fig. 5A). Two dimensions captured stress-resilient phenotypes (1:self-instruction; 2:social and cognitive coping) that loaded high on items from the resilience scale and the coping questionnaire (Fig. 5A). In contrast, two dimensions captured maladaptive stress phenotypes (3:intolerance of uncertainty, 5:avoidance/distraction) that loaded high on the respective IoU and coping subscales. Finally, the fourth dimension (negative affectivity) loaded high on state depressive symptoms and TAI items (Fig. 5B) that are part of the ‘depression’ factor (22). Elevated TAI scores are highly prevalent in mood and anxiety disorders (23), and might have a shared genetic basis (24). Notably, individual scores on the five dimensions of stress adaptation correlated differentially with the subjective response to the psycho-social stress (Supplementary Figure S8 and Table S5).

**Figure 5:**
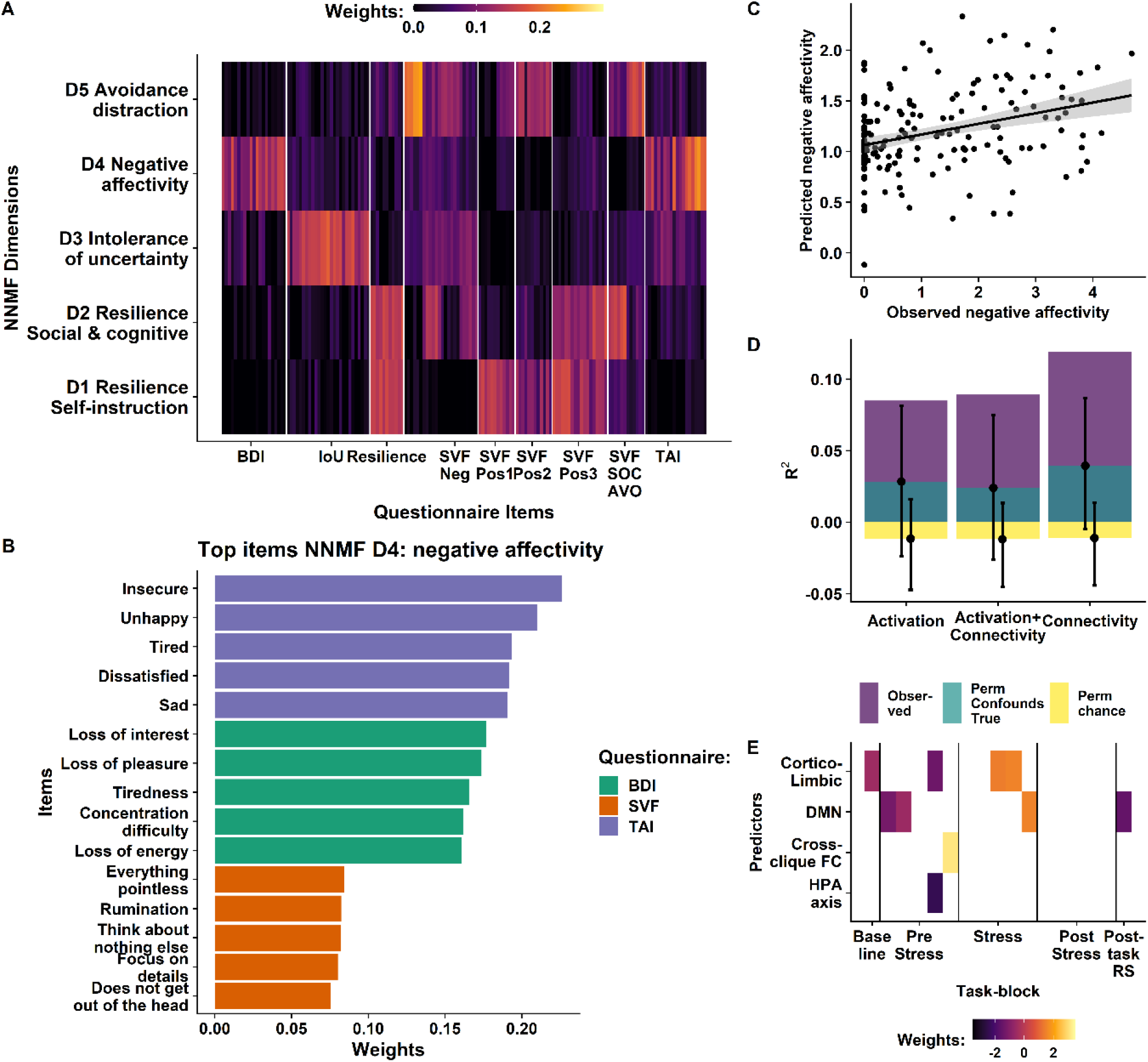
Block-wise changes in functional connectivity (FC) within the four stress subnetworks predict negative affectivity. A: Non-negative matrix factorization (NNMF) revealed 5 dimensions of individual stress responsivity that capture resilient and susceptible phenotypes. B: Weights of representative items contributing to the negative affectivity NNMF dimension. Shown are the top five items from the three questionnaires contributing most items to the dimension. C: A model including stress-induced spatio-temporal FC changes predicts negative affectivity. Predicted and observed values of negative affectivity were significantly correlated (*r*=.35, *p*_perm_=.009) and the model explained 12% variance. D: Adding stress-induced changes in brain responses and FC improves the prediction of negative affectivity compared to permutations of only the response variable (chance level, yellow) or the response variable and the confounding variables age, sex, average framewise displacement and pre-task cortisol response correspondingly (confound baseline, turquoise). Error bars depict 95% percentiles. E: Weights from the combined prediction model including stress-induced changes in brain response and FC (retained weights add to the prediction beyond confounding variables). DMN = default mode network, BDI = Becks depression inventory, TAI = trait anxiety inventory, IoU = intolerance of uncertainty, SVF = coping questionnaire (Stressverarbeitungsfragebogen), SOC = social support, AVO = avoidance.

Next, we evaluated whether these inter-individual differences in stress adaptation can be predicted by stress-induced changes in brain responses and FC using elastic net with nested cross-validation. Individual block-wise changes in FC predicted negative affectivity considerably better than confounding variables alone (*ΔR^2^*=.08; *p_perm_*=.009, Fig. 5C-E). Specifically, lower FC in the anticipatory *PreStress* phase across the DMN and cortico-limbic clusters as well as higher stress-induced FC in the same clusters were associated with higher negative affectivity (Fig. 5E). While block-wise changes in brain responses did slightly improve the prediction compared to confounding variables (*ΔR^2^*=.06; *p_perm_*=.038), combining block-wise changes in brain response and FC changes did not further improve the prediction. Stress-induced changes in FC did not predict other psychological dimensions of stress adaptation, while changes in brain responses predicted resilience: self-instruction (*ΔR^2^*=.091; *p*=.01, Supplementary Figure S10), suggesting that changes in activation and FC contribute to separable psychological constructs.

Since negative affectivity predominantly reflected scores of the BDI and TAI, we used the same algorithm to predict average scores of both questionnaires. Trait anxiety was predicted best based on stress-induced changes in brain responses and FC (combined: *ΔR^2^*=.10; *p_perm_*=.005; activation: *ΔR^2^*=.06; *p_perm_*=.019, FC: *ΔR^2^*=.06; *p_perm_*=.02). In contrast, neither BDI scores nor the presence of a mood or anxiety disorder were predicted better than when we relied only on confounding variables (Supplementary Figure S9-S10).

## 4. Discussion

Symptoms of impaired stress regulation are common across many mental disorders and mapping individual symptoms onto stress-induced brain network reconfigurations may help increase our pathomechanistic understanding of disorders. Here, we characterized dynamic changes in FC and brain response across all phases of a psycho-social stress task in participants with and without a range of mood and anxiety disorders. First, we showed that dynamic stress-induced changes in FC, but not activation, recover the current state of the stress cycle. Second, we showed that only spatio-temporal FC profiles predicted inter-individual differences in a dimension related to negative affectivity, a well-established transdiagnostic marker of heightened stress susceptibility. Third, reduced FC in subnetworks dominated by DMN and cortico-limbic edges during stress anticipation and increased FC during stress added to the prediction, highlighting that anticipatory stress regulation (11,84,85) could help unravel signatures indicative of a key psychopathology dimension of affective disorders (86, 87). Taken together, our results provide a quantitative mapping of dynamic brain connectivity changes in the acute stress response that reflect psychological differences in affective processing (i.e., measures that have been associated with mood and anxiety disorders). Our results highlight the large potential of novel analysis techniques that capitalize on the rich individual information in spatio-temporal brain response profiles to stress, supporting the idea that mood and anxiety disorders can be best understood as disorders arising from individual differences in dynamic network interactions.

Our results derived from the predictive modeling of acute spatio-temporal stress signatures show that dynamic network reconfigurations within the DMN and the cortico-limbic network reflect both stress states and psychopathological risk factors, echoing crucial insights concerning neural signaling dynamics within the stress network (40, 41) (88–90). While previous studies have highlighted characteristic changes in brain responses (46,56,57,91,92), most case-control studies are relatively small and cannot resolve dimensional aspects of psychological stress susceptibility, which may add to the limited convergence of findings across studies (93–96). Thereby, our study adds to the growing concern about heterogeneity (i.e., non-ergodicity) within diagnosis categories (97, 98). By combining dynamic stress-induced changes in brain responses and FC in one predictive model trained in a large transdiagnostic sample, we can capitalize on the rich hierarchical information provided by 217×40min of fMRI timeseries data to derive robust individual markers of stress reactivity. Crucially, our findings are closely in line with recent preclinical work on spatio-temporal signatures of mood and anxiety disorders that reflect state and trait characteristics of stress reactivity (99), indicating large potential for translational approaches (100, 101). Despite our comparably large sample (70, 102), conventional analyses comparing average brain responses between diagnostic groups failed to identify characteristic signatures of stress. In contrast, our spatio-temporal network model recovered both the phases of the stress task as well as a psychopathological dimension of maladaptive stress responses: negative affectivity (103, 104). To conclude, our findings highlight the relevance of unique stress-related network dynamics to better understand psychological responses to acute stress, including susceptibility for mood and anxiety disorders that is of high relevance for translational research or clinical trials.

At a mechanistic level, our findings provide crucial insights into cognitive and affective processes that link altered stress-induced brain function to symptoms of psychopathology at the individual level. In line with recent work on ‘connectomic fingerprints’, individual changes in stress-induced FC showed much higher accuracy in predicting stress states and psychopathology dimension, compared to changes in activation. This demonstrates the large potential of a combined hierarchical model for accurate predictions at an individual level (105, 106), especially during relevant task perturbations (50,105,107,108). Relatedly, it supports the notion that most mental disorders can be conceptualized as network disorders (44), where a dynamic network perspective helps extracting unique information (48, 109) that tracks adaptive responses to stressors more faithfully from a neurobiological perspective (110–112). Specifically, our results suggest that network-based reconfigurations in FC during stress as well as in anticipation of stress, particularly between the DMN and edges of a cortico-limbic subnetwork, are important markers reflecting negative affectivity which is in line with extensive previous research on mood and anxiety disorders (113–115). Notably, neurobiologically-inspired treatments such as TMS target comparable networks to elicit clinically meaningful responses (116) and present-centered psychotherapy has been shown to normalize cortico-limbic processing in stress-related disorders (117). To conclude our predictive model demonstrates that an emphasis on modeling individual differences in psychopathology and network-based reconfiguration within the stress network improves prediction and interpretability (45, 118), dovetailing well with recent insights on effective treatment mechanisms.

Although our study provides an innovative approach to bridge the predictive gap between acute stress reactivity and psychological responsivity, it has limitations that need to be addressed in future work. First, to ensure robust inferences, we had to aggregate connectivity within subnetworks to balance model complexity with the number of participants and larger samples will allow to disentangle the specific contribution of connections between nodes in the future. Second, it is conceivable that stress-induced changes on timescales that are not explicitly modeled with our approach (i.e., specific events such as feedback) could improve the prediction. Third, to establish robustness, replication of the spatio-temporal signatures of negative affectivity in an independent dataset is preferable. Likewise, whether the dynamic changes in network FC generalize to other stress tasks remains to be shown and will be an important step for a better understanding of FC in relation to stress-related disorders. Finally, while previous work has shown that negative affectivity is associated with mood and anxiety disorders, our association between short-term stress-induced FC changes and this psychopathological trait cannot address the question of a causal link.

To summarize, dynamic FC changes within the stress network, but not changes in activation, tracked the current stress phase and corresponding changes in pulse rate. Furthermore, these network-based reconfigurations, driven by reduced cortico-limbic and DMN FC in anticipation of stress and increased FC during stress, predicted the psychopathology dimension negative affectivity. Collectively, our results emphasize that characterizing the neural stress response across the entire cycle by modeling individual signatures with high spatial and temporal resolution in a hierarchical model improves the prediction of key changes within participants (i.e., stress phase, pulse rate) and between participants (i.e., differences in negative affectivity). Crucially, since individual signatures predicted the psychopathology dimension negative affectivity, but not the presence of any mood and anxiety disorder, our study highlights the need for transdiagnostic approaches to better understand the multifaceted psychopathological profile of individuals within broad disorder categories. Therefore, our results offer novel insights into stress-related pathomechanisms of mental disorders, providing a potential endophenotype that may guide future translational research.

## Acknowledgement

We thank Anna Hetzel and Ines Eidner for help with data acquisition and Alina Tontsch, Manfred Uhr and the team of the Max Planck Institute of Psychiatry Biobank for sample processing. NBK received salary support from the University of Tübingen, fortune grant #2453-0-0.

## Author contributions

EBB and PGS were responsible for the study concept and design. MC and PGS validated the paradigm and procedure. AK and NBK conceived the method and AK performed the data analysis. AK wrote the first draft of the manuscript and NBK contributed to the writing. All authors contributed to the interpretation of findings, provided critical revision of the manuscript for important intellectual content and approved the final version for publication.

## References

1. McEwen BS (2003): Mood disorders and allostatic load. Biological Psychiatry 54: 200–207.

2. Aldao A, Nolen-Hoeksema S, Schweizer S (2010): Emotion-regulation strategies across psychopathology: A meta-analytic review. Clinical Psychology Review 30: 217–237.

3. Compas BE, Jaser SS, Bettis AH, Watson KH, Gruhn M, Dunbar JP, et al. (2017): Coping, Emotion Regulation and Psychopathology in Childhood and Adolescence: A Meta-Analysis and Narrative Review. Psychol Bull 143: 939– 991.

4. McEwen BS (2004): Protection and Damage from Acute and Chronic Stress: Allostasis and Allostatic Overload and Relevance to the Pathophysiology of Psychiatric Disorders. Annals of the New York Academy of Sciences 1032: 1– 7.

5. Nolen-Hoeksema S, Wisco BE, Lyubomirsky S (2008): Rethinking Rumination. Perspect Psychol Sci 3: 400–424.

6. de Kloet CS, Vermetten E, Geuze E, Kavelaars A, Heijnen CJ, Westenberg HGM (2006): Assessment of HPA-axis function in posttraumatic stress disorder: Pharmacological and non-pharmacological challenge tests, a review. Journal of Psychiatric Research 40: 550–567.

7. Horowitz MA, Zunszain PA (2015): Neuroimmune and neuroendocrine abnormalities in depression: two sides of the same coin. Annals of the New York Academy of Sciences 1351: 68–79.

8. Mehta D, Binder EB (2012): Gene × environment vulnerability factors for PTSD: the HPA-axis. Neuropharmacology 62: 654–662.

9. Zorn JV, Schür RR, Boks MP, Kahn RS, Joëls M, Vinkers CH (2017): Cortisol stress reactivity across psychiatric disorders: A systematic review and meta-analysis. Psychoneuroendocrinology 77: 25–36.

10. Schiweck C, Piette D, Berckmans D, Claes S, Vrieze E (2019): Heart rate and high frequency heart rate variability during stress as biomarker for clinical depression. A systematic review. Psychol Med 49: 200–211.

11. Gaab J, Rohleder N, Nater UM, Ehlert U (2005): Psychological determinants of the cortisol stress response: the role of anticipatory cognitive appraisal. Psychoneuroendocrinology 30: 599–610.

12. Pulopulos MM, Baeken C, De Raedt R (2020): Cortisol response to stress: The role of expectancy and anticipatory stress regulation. Hormones and Behavior 117: 104587.

13. Selye H (1976): Stress without distress. Psychopathology of Human Adaptation. Springer, pp 137–146.

14. de Kloet ER, Joëls M (2020): Mineralocorticoid Receptors and Glucocorticoid Receptors in HPA Stress Responses During Coping and Adaptation. In: de Kloet ER, Joëls M. Oxford Research Encyclopedia of Neuroscience. Oxford University Press. https://doi.org/10.1093/acrefore/9780190264086.013.266

15. Brosschot JF, Gerin W, Thayer JF (2006): The perseverative cognition hypothesis: A review of worry, prolonged stress-related physiological activation, and health. Journal of Psychosomatic Research 60: 113–124.

16. Gold SM, Zakowski SG, Valdimarsdottir HB, Bovbjerg DH (2004): Higher Beck depression scores predict delayed epinephrine recovery after acute psychological stress independent of baseline levels of stress and mood. Biological Psychology 67: 261–273.

17. Lü W, Wang Z, You X (2016): Physiological responses to repeated stress in individuals with high and low trait resilience. Biological Psychology 120: 46–52.

18. Morris MC, Kouros CD, Mielock AS, Rao U (2017): Depressive Symptom Composites Associated with Cortisol Stress Reactivity in Adolescents. J Affect Disord 210: 181–188.

19. Morris MC, Rao U, Garber J (2012): Cortisol responses to psychosocial stress predict depression trajectories: Social-evaluative threat and prior depressive episodes as moderators. Journal of Affective Disorders 143: 223–230.

20. Rudolph KD, Troop-Gordon W, Granger DA (2011): Individual differences in biological stress responses moderate the contribution of early peer victimization to subsequent depressive symptoms. Psychopharmacology 214: 209–219.

21. Fiksdal A, Hanlin L, Kuras Y, Gianferante D, Chen X, Thoma MV, Rohleder N (2019): Associations between symptoms of depression and anxiety and cortisol responses to and recovery from acute stress. Psychoneuroendocrinology 102: 44–52.

22. Balsamo M, Romanelli R, Innamorati M, Ciccarese G, Carlucci L, Saggino A (2013): The State-Trait Anxiety Inventory: Shadows and Lights on its Construct Validity. J Psychopathol Behav Assess 35: 475–486.

23. Knowles KA, Olatunji BO (2020): Specificity of trait anxiety in anxiety and depression: Meta-analysis of the State-Trait Anxiety Inventory. Clinical Psychology Review 82: 101928.

24. Thorp JG, Campos AI, Grotzinger AD, Gerring ZF, An J, Ong J-S, et al. (2021): Symptom-level modelling unravels the shared genetic architecture of anxiety and depression. Nat Hum Behav 1–11.

25. Gecaite J, Burkauskas J, Brozaitiene J, Mickuviene N (2019): Cardiovascular Reactivity to Acute Mental Stress: The Importance of Type D Personality, Trait Anxiety, and Depression Symptoms in Patiens after Acute Coronary Syndromes. Journal of Cardiopulmonary Rehabilitation and Prevention 39: E12.

26. Quirin M, Kazén M, Rohrmann S, Kuhl J (2009): Implicit but Not Explicit Affectivity Predicts Circadian and Reactive Cortisol: Using the Implicit Positive and Negative Affect Test. Journal of Personality 77: 401–426.

27. Zellars KL, Meurs JA, Perrewé PL, Kacmar CJ, Rossi AM (2009): Reacting to and recovering from a stressful situation: The negative affectivity-physiological arousal relationship. Journal of Occupational Health Psychology 14: 11–22.

28. LeMoult J (2020): From Stress to Depression: Bringing Together Cognitive and Biological Science. Curr Dir Psychol Sci 29: 592–598.

29. Schlotz W, Yim IS, Zoccola PM, Jansen L, Schulz P (2011): The perceived stress reactivity scale: Measurement invariance, stability, and validity in three countries. Psychological Assessment 23: 80–94.

30. LeMoult J, Arditte KA, D’Avanzato C, Joormann J (2013): State Rumination: Associations with Emotional Stress Reactivity and Attention Biases. Journal of Experimental Psychopathology 4: 471–484.

31. Ottaviani C, Thayer JF, Verkuil B, Lonigro A, Medea B, Couyoumdjian A, Brosschot JF (2016): Physiological concomitants of perseverative cognition: A systematic review and meta-analysis. Psychological Bulletin 142: 231–259.

32. Quinn ME, Grant KE, Adam EK (2018): Negative cognitive style and cortisol recovery accentuate the relationship between life stress and depressive symptoms. Stress 21: 119–127.

33. Stewart JG, Mazurka R, Bond L, Wynne-Edwards KE, Harkness KL (2013): Rumination and Impaired Cortisol Recovery Following a Social Stressor in Adolescent Depression. J Abnorm Child Psychol 41: 1015–1026.

34. Janson J, Rohleder N (2017): Distraction coping predicts better cortisol recovery after acute psychosocial stress. Biological Psychology 128: 117–124.

35. Salzmann S, Euteneuer F, Strahler J, Laferton JAC, Nater UM, Rief W (2018): Optimizing expectations and distraction leads to lower cortisol levels after acute stress. Psychoneuroendocrinology 88: 144–152.

36. Meuwly N, Bodenmann G, Germann J, Bradbury TN, Ditzen B, Heinrichs M (2012): Dyadic coping, insecure attachment, and cortisol stress recovery following experimentally induced stress. Journal of Family Psychology 26: 937–947.

37. Raymond C, Marin M-F, Juster R-P, Lupien SJ (2019): Should we suppress or reappraise our stress?: the moderating role of reappraisal on cortisol reactivity and recovery in healthy adults. *Anxiety, Stress*, & Coping 32: 286–297.

38. Cheetham-Blake TJ, Turner-Cobb JM, Family HE, Turner JE (2019): Resilience characteristics and prior life stress determine anticipatory response to acute social stress in children aged 7–11 years. British Journal of Health Psychology 24: 282–297.

39. Mikolajczak M, Roy E, Luminet O, Timary P de (2008): Resilience and hypothalamic-pituitary-adrenal axis reactivity under acute stress in young men. Stress 11: 477–482.

40. Hermans EJ, Henckens MJAG, Joëls M, Fernández G (2014): Dynamic adaptation of large-scale brain networks in response to acute stressors. Trends Neurosci 37: 304–314.

41. van Oort J, Tendolkar I, Hermans EJ, Mulders PC, Beckmann CF, Schene AH, et al. (2017): How the brain connects in response to acute stress: A review at the human brain systems level. Neuroscience & Biobehavioral Reviews 83: 281– 297.

42. Vaisvaser S, Lin T, Admon R, Podlipsky I, Greenman Y, Stern N, et al. (2013): Neural traces of stress: cortisol related sustained enhancement of amygdala-hippocampal functional connectivity. Front Hum Neurosci 7. https://doi.org/10.3389/fnhum.2013.00313

43. Zhang W, Llera A, Hashemi MM, Kaldewaij R, Koch SBJ, Beckmann CF, et al. (2020): Discriminating stress from rest based on resting-state connectivity of the human brain: A supervised machine learning study. Human Brain Mapping n/a. https://doi.org/10.1002/hbm.25000

44. McTeague LM, Rosenberg BM, Lopez JW, Carreon DM, Huemer J, Jiang Y, et al. (2020): Identification of Common Neural Circuit Disruptions in Emotional Processing Across Psychiatric Disorders. Am J Psychiatry 177: 411–421.

45. Corr R, Pelletier-Baldelli A, Glier S, Bizzell J, Campbell A, Belger A (2020): Neural Mechanisms of Acute Stress and Trait Anxiety in Adolescents. NeuroImage: Clinical 102543.

46. Wheelock MD, Harnett NG, Wood KH, Orem TR, Granger DA, Mrug S, Knight DC (2016): Prefrontal Cortex Activity Is Associated with Biobehavioral Components of the Stress Response. Front Hum Neurosci 10. https://doi.org/10.3389/fnhum.2016.00583

47. Braun U, Schäfer A, Bassett DS, Rausch F, Schweiger JI, Bilek E, et al. (2016): Dynamic brain network reconfiguration as a potential schizophrenia genetic risk mechanism modulated by NMDA receptor function. PNAS 113: 12568–12573.

48. Braun U, Schaefer A, Betzel RF, Tost H, Meyer-Lindenberg A, Bassett DS (2018): From Maps to Multi-dimensional Network Mechanisms of Mental Disorders. Neuron 97: 14–31.

49. Finn ES, Todd Constable R (2016): Individual variation in functional brain connectivity: implications for personalized approaches to psychiatric disease. Dialogues Clin Neurosci 18: 277–287.

50. Greene AS, Gao S, Noble S, Scheinost D, Constable RT (2020): How Tasks Change Whole-Brain Functional Organization to Reveal Brain-Phenotype Relationships. Cell Reports 32: 108066.

51. Figee M, Mayberg H (2021): The future of personalized brain stimulation. Nat Med 27: 196–197.

52. Brückl TM, Spoormaker VI, Sämann PG, Brem A-K, Henco L, Czamara D, et al. (2020): The biological classification of mental disorders (BeCOME) study: a protocol for an observational deep-phenotyping study for the identification of biological subtypes. BMC Psychiatry 20: 213.

53. Wittchen H, Beloch E, Garczynski E, Holly A, Lachner G, Perkonigg A, et al. (1995): Münchener Composite International Diagnostic Interview (M-CIDI). *München*: *Max-Planck-Institut für Psychiatrie, Klinisches Institut*.

54. Muehlhan M, Lueken U, Wittchen H-U, Kirschbaum C (2011): The scanner as a stressor: Evidence from subjective and neuroendocrine stress parameters in the time course of a functional magnetic resonance imaging session. International Journal of Psychophysiology 79: 118–126.

55. Janke W (1994): Befindlichkeitsskalierung durch Kategorien und Eigenschaftswörter: BSKE (EWL) nach Janke, Debus, Erdmann und Hüppe. Test und Handanweisung Unveröffentlichter Institutsbericht, Lehrstuhl für Biologische und Klinische Psychologie der Universität Würzburg.

56. Pruessner JC, Dedovic K, Khalili-Mahani N, Engert V, Pruessner M, Buss C, et al. (2008): Deactivation of the Limbic System During Acute Psychosocial Stress: Evidence from Positron Emission Tomography and Functional Magnetic Resonance Imaging Studies. Biological Psychiatry 63: 234–240.

57. Elbau IG, Brücklmeier B, Uhr M, Arloth J, Czamara D, Spoormaker VI, et al. (2018): The brain’s hemodynamic response function rapidly changes under acute psychosocial stress in association with genetic and endocrine stress response markers. PNAS 201804340.

58. Kühnel A, Kroemer NB, Elbau IG, Czisch M, Sämann PG, Walter M, Binder EB (2020): Psychosocial stress reactivity habituates following acute physiological stress. Human Brain Mapping 41: 4010–4023.

59. Beck AT, Steer RA, Brown G (1996): Beck depression inventory–II. Psychological Assessment.

60. Spielberger C, Gorsuch R, Lushene R, Vagg P, Jacobs G (1983): Manual for the State-Trait Anxiety Inventory (Form Y) Mind Garden. Palo Alto, CA.

61. Gerlach AL, Andor T, Patzelt J (2008): The significance of intolerance of uncertainty in generalized anxiety disorder: Possible models and development of a German version of the intolerance of uncertainty scale. ZEITSCHRIFT FUR KLINISCHE PSYCHOLOGIE UND PSYCHOTHERAPIE 37: 190–199.

62. Ising M, Weyers P, Janke W, Erdmann G (2001): The psychometric properties of the SVF78 by Janke and Erdmann, a short version of the SVF120. Zeitschrift fur Differentielle und Diagnostische Psychologie 22: 279–290.

63. Wagnild GM, Young HM (1993): Development and Psychometric Evaluation of the Resilience Scale. Journal of nursing measurement 1: 165–17847.

64. Ashburner J (2007): A fast diffeomorphic image registration algorithm. NeuroImage 38: 95–113.

65. Lee DD, Seung HS (1999): Learning the parts of objects by non-negative matrix factorization [no. 6755]. Nature 401: 788–791.

66. Cattell RB (1966): The scree test for the number of factors. Multivariate behavioral research 1: 245–276.

67. Wust S, Wolf J, Hellhammer DH, Federenko I, Schommer N, Kirschbaum C (2000): The cortisol awakening response - normal values and confounds. Noise and Health 2: 79.

68. Finn ES, Shen X, Scheinost D, Rosenberg MD, Huang J, Chun MM, et al. (2015): Functional connectome fingerprinting: Identifying individuals based on patterns of brain connectivity. Nat Neurosci 18: 1664–1671.

69. Shen X, Tokoglu F, Papademetris X, Constable RT (2013): Groupwise whole-brain parcellation from resting-state fMRI data for network node identification. Neuroimage 0: 403–415.

70. Noack H, Nolte L, Nieratschker V, Habel U, Derntl B (2019): Imaging stress: an overview of stress induction methods in the MR scanner. J Neural Transm 126: 1187–1202.

71. Behzadi Y, Restom K, Liau J, Liu TT (2007): A component based noise correction method (CompCor) for BOLD and perfusion based fMRI. NeuroImage 37: 90– 101.

72. Kroemer NB, Sun X, Veldhuizen MG, Babbs AE, de Araujo IE, Small DM (2016): Weighing the evidence: Variance in brain responses to milkshake receipt is predictive of eating behavior. NeuroImage 128: 273–283.

73. Kroemer NB, Guevara A, Ciocanea Teodorescu I, Wuttig F, Kobiella A, Smolka MN (2014): Balancing reward and work: Anticipatory brain activation in NAcc and VTA predict effort differentially. NeuroImage 102: 510–519.

74. McLaren DG, Ries ML, Xu G, Johnson SC (2012): A Generalized Form of Context-Dependent Psychophysiological Interactions (gPPI): A Comparison to Standard Approaches. Neuroimage 61: 1277–1286.

75. Mejia AF, Nebel MB, Barber AD, Choe AS, Pekar JJ, Caffo BS, Lindquist MA (2018): Improved estimation of subject-level functional connectivity using full and partial correlation with empirical Bayes shrinkage. Neuroimage 172: 478– 491.

76. Narayan M, Allen GI (2016): Mixed Effects Models for Resampled Network Statistics Improves Statistical Power to Find Differences in Multi-Subject Functional Connectivity. Front Neurosci 10. https://doi.org/10.3389/fnins.2016.00108

77. Pervaiz U, Vidaurre D, Woolrich MW, Smith SM (2020): Optimising network modelling methods for fMRI. NeuroImage 211: 116604.

78. Charrad M, Ghazzali N, Boiteau V, Niknafs A, Charrad MM (2014): Package ‘nbclust.’ Journal of statistical software 61: 1–36.

79. Pedregosa F, Varoquaux G, Gramfort A, Michel V, Thirion B, Grisel O, et al. (2011): Scikit-learn: Machine Learning in Python. Journal of Machine Learning Research 12: 2825–2830.

80. Wager TD, Atlas LY, Lindquist MA, Roy M, Woo C-W, Kross E (2013): An fMRI-Based Neurologic Signature of Physical Pain. New England Journal of Medicine 368: 1388–1397.

81. Jollans L, Boyle R, Artiges E, Banaschewski T, Desrivières S, Grigis A, et al. (2019): Quantifying performance of machine learning methods for neuroimaging data. NeuroImage 199: 351–365.

82. R Core Team (2018): R: A Language and Environment for Statistical Computing. Vienna, Austria: R Foundation for Statistical Computing. Retrieved from https://www.R-project.org/

83. Kuznetsova A, Brockhoff P, Christensen R (2017): lmerTest Package: Tests in Linear Mixed Effects Models. https://doi.org/10.18637/JSS.V082.I13

84. Fehlner P, Bilek E, Harneit A, Böhringer A, Moessnang C, Meyer-Lindenberg A, Tost H (2020): Neural responses to social evaluative threat in the absence of negative investigator feedback and provoked performance failures. Human Brain Mapping 41: 2092–2103.

85. McKlveen JM, Myers B, Herman JP (2015): The Medial Prefrontal Cortex: Coordinator of Autonomic, Neuroendocrine and Behavioural Responses to Stress. J Neuroendocrinol 27: 446–456.

86. Böhnke JR, Lutz W, Delgadillo J (2014): Negative affectivity as a transdiagnostic factor in patients with common mental disorders. Journal of Affective Disorders 166: 270–278.

87. Hur J, Stockbridge MD, Fox AS, Shackman AJ (2019): Chapter 16 - Dispositional negativity, cognition, and anxiety disorders: An integrative translational neuroscience framework. In: Srinivasan N, editor. Progress in Brain Research, vol. 247. Elsevier, pp 375–436.

88. Goldfarb EV, Rosenberg MD, Seo D, Constable RT, Sinha R (2020): Hippocampal seed connectome-based modeling predicts the feeling of stress [no. 1]. Nature Communications 11: 2650.

89. Wheelock MD, Harnett NG, Wood KH, Orem TR, Mrug S, Deshpande G, et al. (n.d.): Psychosocial Stress Reactivity Is Associated With Decreased Whole-Brain Network Efficiency and Increased Amygdala Centrality. 13.

90. Zhang W, Hashemi MM, Kaldewaij R, Koch SBJ, Beckmann C, Klumpers F, Roelofs K (2019): Acute stress alters the ‘default’ brain processing. NeuroImage 189: 870–877.

91. Dedovic K, D’Aguiar C, Pruessner JC (2009): What Stress Does to Your Brain: A Review of Neuroimaging Studies. The Canadian Journal of Psychiatry 54: 6– 15.

92. Lederbogen F, Ulshöfer E, Peifer A, Fehlner P, Bilek E, Streit F, et al. (2018): No association between cardiometabolic risk and neural reactivity to acute psychosocial stress. NeuroImage: Clinical 20: 1115–1122.

93. Admon R, Holsen LM, Aizley H, Remington A, Whitfield-Gabrieli S, Goldstein JM, Pizzagalli DA (2015): Striatal hyper-sensitivity during stress in remitted individuals with recurrent depression. Biol Psychiatry 78: 67–76.

94. Oort J van, Kohn N, Vrijsen JN, Collard R, Duyser FA, Brolsma SCA, et al. (2020): Absence of default mode downregulation in response to a mild psychological stressor marks stress-vulnerability across diverse psychiatric disorders. NeuroImage: Clinical 102176.

95. Villarreal MF, Wainsztein AE, Mercè RÁ, Goldberg X, Castro MN, Brusco LI, et al. (2021): Distinct Neural Processing of Acute Stress in Major Depression and Borderline Personality Disorder. Journal of Affective Disorders S0165032721001890.

96. Waugh CE, Hamilton JP, Chen MC, Joormann J, Gotlib IH (2012): Neural temporal dynamics of stress in comorbid major depressive disorder and social anxiety disorder. Biology of Mood & Anxiety Disorders 2: 11.

97. Adolf JK, Fried EI (2019): Ergodicity is sufficient but not necessary for group-to-individual generalizability. PNAS 116: 6540–6541.

98. Fisher AJ, Medaglia JD, Jeronimus BF (2018): Lack of group-to-individual generalizability is a threat to human subjects research. Proc Natl Acad Sci USA 115: E6106–E6115.

99. Hultman R, Ulrich K, Sachs BD, Blount C, Carlson DE, Ndubuizu N, et al. (2018): Brain-wide Electrical Spatiotemporal Dynamics Encode Depression Vulnerability. Cell 173: 166–180.e14.

100. Flores Á, Fullana MÀ, Soriano-Mas C, Andero R (2018): Lost in translation: how to upgrade fear memory research [no. 11]. Molecular Psychiatry 23: 2122–2132.

101. Notaras M, van den Buuse M (2020): Neurobiology of BDNF in fear memory, sensitivity to stress, and stress-related disorders [no. 10]. Molecular Psychiatry 25: 2251–2274.

102. Kogler L, Mueller VI, Chang A, Eickhoff SB, Fox PT, Gur RC, Derntl B (2015): Psychosocial versus physiological stress – meta-analyses on deactivations and activations of the neural correlates of stress reactions. Neuroimage 119: 235– 251.

103. Gulley LD, Hankin BL, Young JF (2016): Risk for Depression and Anxiety in Youth: The Interaction between Negative Affectivity, Effortful Control, and Stressors. J Abnorm Child Psychol 44: 207–218.

104. Hur J, Kuhn M, Grogans SE, Anderson AS, Islam S, Kim HC, et al. (2021): Anxiety-related frontocortical activity is associated with dampened stressor reactivity in the real world. bioRxiv 2021.03.17.435791.

105. Finn ES, Scheinost D, Finn DM, Shen X, Papademetris X, Constable RT (2017): Can brain state be manipulated to emphasize individual differences in functional connectivity? NeuroImage 160: 140–151.

106. Waller L, Walter H, Kruschwitz JD, Reuter L, Müller S, Erk S, Veer IM (2017): Evaluating the replicability, specificity, and generalizability of connectome fingerprints. NeuroImage 158: 371–377.

107. Cole MW, Ito T, Bassett DS, Schultz DH (2016): Activity flow over resting-state networks shapes cognitive task activations [no. 12]. Nature Neuroscience 19: 1718–1726.

108. Cole MW, Bassett DS, Power JD, Braver TS, Petersen SE (2014): Intrinsic and Task-Evoked Network Architectures of the Human Brain. Neuron 83: 238–251.

109. Fornito A, Zalesky A, Breakspear M (2015): The connectomics of brain disorders [no. 3]. Nature Reviews Neuroscience 16: 159–172.

110. Alexander L, Wood CM, Gaskin PLR, Sawiak SJ, Fryer TD, Hong YT, et al. (2020): Over-activation of primate subgenual cingulate cortex enhances the cardiovascular, behavioral and neural responses to threat [no. 1]. Nature Communications 11: 5386.

111. Grueschow M, Stenz N, Thörn H, Ehlert U, Breckwoldt J, Brodmann Maeder M, et al. (2021): Real-world stress resilience is associated with the responsivity of the locus coeruleus [no. 1]. Nature Communications 12: 2275.

112. Sousa N (2016): The dynamics of the stress neuromatrix [no. 3]. Molecular Psychiatry 21: 302–312.

113. Sikora M, Heffernan J, Avery ET, Mickey BJ, Zubieta J-K, Peciña M (2016): Salience Network Functional Connectivity Predicts Placebo Effects in Major Depression. Biological Psychiatry: Cognitive Neuroscience and Neuroimaging 1: 68–76.

114. Sripada RK, King AP, Welsh RC, Garfinkel SN, Wang X, Sripada CS, Liberzon I (2012): Neural Dysregulation in Posttraumatic Stress Disorder: Evidence for Disrupted Equilibrium between Salience and Default Mode Brain Networks. Psychosom Med 74: 904–911.

115. Whitton AE, Webb CA, Dillon DG, Kayser J, Rutherford A, Goer F, et al. (2019): Pretreatment Rostral Anterior Cingulate Cortex Connectivity With Salience Network Predicts Depression Recovery: Findings From the EMBARC Randomized Clinical Trial. Biological Psychiatry 85: 872–880.

116. Philip NS, Barredo J, van ’t Wout-Frank M, Tyrka AR, Price LH, Carpenter LL (2018): Network Mechanisms of Clinical Response to Transcranial Magnetic Stimulation in Posttraumatic Stress Disorder and Major Depressive Disorder. Biological Psychiatry 83: 263–272.

117. Abdallah CG, Averill CL, Ramage AE, Averill LA, Alkin E, Nemati S, et al. (2019): Reduced Salience and Enhanced Central Executive Connectivity Following PTSD Treatment. Chronic Stress 3: 2470547019838971.

118. Sinha R, Lacadie CM, Constable RT, Seo D (2016): Dynamic neural activity during stress signals resilient coping. Proc Natl Acad Sci USA 113: 8837–8842.

