## Supplementary Information for "Spatio-temporal dynamics of stress-induced network reconfigurations reflect negative affectivity"

[#] contributed equally

**Corresponding authors***

**Assessment of subjective stress experience (BSKE scales):**

The BSKE (*Befindlichkeitsskalierung durch Kategorien und Eigenschaftswörter*, Janke, 1994) scales are a short version of the more extensive "*Eigenschaftswörterliste*" (EWL, (Janke and Debus, 1978), a scale developed to assess current emotional state across positive and negative dimensions. The scale consists of 15 items (emotions / states) and participants were asked to rate their current state/feeling ("I feel …") on 6-point scale ranging from 1 ("not at all / gar nicht") to 6 ("very strongly / sehr stark"). We calculated sum scores including the items activity, wakefulness, self-certainty, focus, and relaxed state of mind for positive affect and including the items internal and external agitation, anxiety, sadness, anger, dysphoria, sensitivity as well as three items assessing somatic changes for negative affect.

**Procedure:**

On the day, participants arrived at the scanner at approximately 10 am. Upon arrival, the first saliva sample was taken to measure basal cortisol levels and an intravenous catheter was placed for blood sampling in a subsample (N=73). After that, participants were familiarized with the task and the response options. Electrodes were placed on the palm of the left hand for the measurement of skin conductance and on the back for electrocardiography. A pulse oximeter was placed on the fingertip to measure pulse rate. Before entering the scanner, we took another saliva sample. The fMRI session started with a T2-weighted high-resolution image for spatial normalization, followed by an emotional face matching task and a pre-stress resting state. Subsequently, participants completed the psychosocial stress task. At the end participants rated their subjective state again and another saliva sample was taken. After a 30min break outside of the scanner a last resting state fMRI was acquired followed by another saliva sample and subjective state ratings.

**Paradigm:**

The psychosocial stress-task was the same as used and described in a previous publication (Kühnel et al., 2020). Participants had to solve mental arithmetic problems either in a control condition without time pressure and negative feedback or under stress with a time limit and negative feedback. Critically, the task had three phases, *PreStress*, *Stress*, and *PostStress*, each consisting of five 50 second blocks of arithmetic each followed by 40 second rest blocks (fixation cross). During an arithmetic block, participants were presented an arithmetic problem with a solution between 0 and 9. Arithmetic problems varied in their difficulty across three levels and difficulty was balanced across the three phases. The correct answer was chosen using a response box allowing to navigate a two-button dial wheel system. After selecting the answer, the screen 'froze' for an anticipation phase (2.5 ± 1s, jittered) that was followed by the feedback ('correct', 'incorrect' or 'timeout', presented for 660ms). During *PreStress* and *PostStress*, participants had 10.5 seconds to solve the problem and respond and no further evaluative feedback or cues were given. Before *Stress*, participants were informed that answers are now 'recorded'. During stress, time to solve the arithmetic was generally limited to 4.5 seconds, and in part self-adaptive depending on the participant's preceding performance (i.e., response time was shortened if the participant performed well). Further, a time bar indicated how much time was left, inducing further time pressure, and a performance indicator showed that current performance was below group average ('in the red area'). Two instances of scripted negative verbal feedback in two rest periods informed the participants about their sub-par performance and pushed them to work harder.

#### Physiological recording and preprocessing:

As described in Kühnel et al. (2020), the PPG data was acquired with an MR compatible pulse oximeter (Nonin Medical Inc., Plymouth MN, USA) attached to the pulp of the left ring finger. PPG data, sampled at 5 kHz, was amplified using a MR compatible multi-channel BrainVision ExG AUX Box coupled with a BrainVision ExG MR Amplifier (Brain Products GmbH, Gilching, Germany) and recorded with BrainVision Recorder software 1.0. After down-sampling to 100 Hz, RR-intervals were detected using the Physionet Cardiovascular Signal toolbox (Vest et al., 2018). Success of detection of beat positions was evaluated by visual inspection. Measurements with insufficient data-quality leading to failed detection of beat positions were excluded (n= 27). Subsequent analysis of the pulse rate was based on the derived RR-intervals and conducted with the RHRV package (Rodríguez-Liñares et al., 2008) for R. Further preprocessing involved the exclusion of implausible interbeat-intervals (IBI). We filtered out IBIs shorter than 0.3 s and longer than 2.4s and excluded IBIs showing excessive deviations from the previous, following, or running average (50 beats) IBI. The threshold for excessive deviations was updated dynamically with the initial threshold set at 13% change from IBI to IBI (Vila et al., 1997).

#### Saliva cortisol measurement:

Cortisol concentrations were measured repeatedly before, during, and after the task in saliva and/or serum (Figure 1). Salivary cortisol was sampled directly at arrival before placement of the intravenous catheter for blood sampling in a subsample (T1), 20 minutes later before entering the MRI Scanner (T2) and directly after the stress paradigm (T6) and 30 minutes after the end (T8). After collection, all probes were centrifuged and stored at -80° C until further processing. Salivary cortisol concentrations were measured with electro-chemiluminescence-assay (ECLIA) kit (Cobas®, Roche Diagnostics GmbH, Mannheim, Germany). The detection limit was 1090 pg/mL. The %CV (coefficient of variation) in saliva samples with varying concentrations was between 2.5% and 6.1% for intra-assay variability and between 3.6% und 11.8% for inter-assay variability.

#### fMRI Imaging parameters:

The following scanner settings were used for acquisition of echo-planar images (EPI) for the imaging stress task: 40 oblique slices, oriented along the AC-PC plane, covering the whole brain, interleaved ascending acquisition order, TR= 2s, TE= 40ms, 64 × 64 matrix, field of view = 200 × 200 $mm^2$, voxel size = 3.5 × 3.5 × 3 $mm^3$. The EPI for the two resting state measurements (fixation cross, eyes open) were acquired with the following parameters: 42 oblique slices, oriented along the AC-PC plane, covering the whole brain, interleaved ascending acquisition order, TR= 2.5s, TE= 30ms, 96 × 96 matrix, field of view = 240 × 240 $mm^2$, voxel size = 3.5 × 3.5 × 3 $mm^3$. Additionally, the measurements included a structural high resolution T1 image and single spin-echo EPI volume with the same geometrical settings as the fMRI sequence, but a longer TR of 10 s and TE 0f 38.4 ms. This single EPI volume has the same geometric distortions as the fMRI images, as the k-space readout was identical. Yet, the image combines a higher signal-to-noise ratio and less susceptibility induced signal drop-out, and was used for normalization to correct for field distortions.

#### fMRI preprocessing and movement covariates:

We used the same preprocessing pipeline as reported in Kühnel et al (2020) for both the task and resting state fMRI data. First, data was slice-time corrected and realigned to the first image of the task to correct for head motion. For spatial normalization a single high-resolution spin-echo EPI image was segmented using the unified segmentation scheme. Extracted gray matter and white matter segments were used for DARTEL (Ashburner, 2007) normalization to MNI templates. Functional images were co-registered to the single EPI image and normalized by applying the DARTEL-derived transformation matrix. Data was interpolated with a resolution of 2x2x2 $mm^3$. The last step was the smoothing of the data with a 6x6x6 $mm^3$ full width at half maximum kernel. During the realignment, the six head motion-parameters were extracted for later use as nuisance covariates. Additionally, we extracted physiological noise components based on aCompCor (Behzadi et al., 2007). We extracted the voxel-wise timeseries of the normalized but unsmoothed functional data from thresholded (p> .90) white matter and cerebro-spinal fluid segments, performed PCA, and used the first five components of each segment as physiological noise covariates. Moreover, we calculated the average framewise displacement for all six head motion parameters (FD) and DVARS (spatial root mean square of the data after temporal differencing) for each task-block as well as for the complete task run for later use as a person-level confounding variable capturing interindividual differences in head motion. On the block level, those movement variables differed between task-phases. Nonetheless, a prediction of the task-phase based on the average FD and DVARS of each task-block using the same support-vector machine algorithm as for the FC based data only reached an accuracy of 40% well below the accuracy of the FC based prediction and performed better in only 9 out of 221 participants (Figure S10). Moreover, individual predictive performance of the support vector machine based either on FD and DVARS or FC changes was only weakly correlated ($r$ = .17). Since on the participant-level FD and DVARS were highly correlated ($r$ = .89, $p$ < .001) and neither was correlated with the outcomes of interest (i.e., psychometric dimensions of stress reactivity and diagnosis status) we only included FD as a confounding variable in the elastic-net prediction. In line with previous studies (Goldfarb et al., 2019) we would have excluded runs exceeding an average framewise displacement > 1.5 mm. However, all runs fulfilled this criterion and were included.

#### fMRI First-level design and similarity:

The first-level general linear models (GLM) were built using individual onsets and durations of all task-blocks extracted from the log files for each participant. The task was modelled with three regressors, each modeling the five arithmetic blocks (50s) from the conditions *PreStress*, *Stress* and *PostStress*, respectively. In addition, we included two regressors modeling individual motor responses and verbal feedback during the Stress phase. Nuisance regressors were the six movement parameters derived from realignment, their derivatives, and five physiological noise components extracted from white matter and cerebro-spinal fluid each. Data were high pass filtered with a cut-off of 256s. The contrasts of interest, Stress – *PreStress*, to assess acute psychsocial stress, and *PostStress* – *PreStress*, to assess effects of fast stress recovery, were estimated for each participant.

To capture individual neural stress effects independent of their directionality and localization, we calculated within-participant similarity of the neural activity during *PreStress* compared to *Stress* and *PostStress*, respectively. To this end, we extracted mean beta estimates of the conditions

(*PreStress*, *Stress*, and *PostStress*) from 268 regions of interest (ROIs) spanning the whole brain using an established brain parcellation (Shen et al., 2013). Representational similarity was then calculated as the z-standardized Pearson correlation between the activity *PreStress* and *Stress* or PressStress and *PostStress* across all ROIs for each participant separately.

##### Concatenation of resting-state and task timeseries:

To assess task-induced functional connectivity changes referenced to a resting-state baseline we concatenated timeseries data from the psycho-social stress task and the two, flanking resting-states. Timeseries were linearly detrended (we did not include a quadratic trend to prevent excluding potential task effects with the same pattern induced by the task structure with non-stress phases flanking the acute stress), despiked, and denoised for each measurement separately so that the average gray scale values of each measurement was 0. To concatenate the task timeseries with the resting states, we matched the average gray scale values of the flanking resting-states with the average gray scale value of the rest baseline phases (fixation cross) during the *PreStress* condition for each region of interest. To this end, we calculated the average gray scale value for each region of interest of the rest baseline phases (fixation cross) during the *PreStress* condition and then subtracted this offset from the complete task timeseries, so that the average gray scale value during the rest baseline phases during *PreStress* was 0 and matched the average gray scale values of the flanking resting-states.

##### Prediction of pulse rate changes within conditions and interindividual differences:

To test whether the predictive performance was only driven by the differences between difference between the stress and non-stress conditions we applied linear-mixed effects models within conditions. Predicted and observed changes in pulse rate were significantly associated in the *PostStress* ($b = .11$, $p < .001$) and to a lesser extent Stress condition ($b = .07$, $p < .001$) and *PreStress* ($b = .03$, $p = .02$) task-blocks (Fig. 4D), indicating that especially FC-changes during stress recovery correspond to the recovery in the autonomous response. Moreover, interindividual differences in stress-induced pulse rate changes (i.e. *Stress* – *PreStress*) were correlated (*Stress*: $r=.22$, $p = .002$, *PostStress*: $r = .25$, $p = .0004$ Fig. 4E-F) when deriving them either from observed or predicted pulse rate changes.

##### General Psychopathology factor (P-factor):

In recent years a general psychopathology factor (p-factor) has become more popular to assess the overall strength of psychopathology for each individual across the complete spectrum of disorders (Caspi et al., 2014). Since p factors derived using factor analytic approaches on single symptoms have been shown to correlate highly with simple additive operationalizations of the p-factor (i.e., adding the number of symptoms / diagnoses, (Fried et al., 2021)), we derived a p-factor score by adding all subthreshold (1) and full diagnoses (2) within the last 12 months derived from the CIDI interview. While this p-factor was correlated with the positive and negative subjective response to the stressor as well as the non-negative matrix factorization dimensions negative affectivity and intolerance of uncertainty (Fig. S8), we could not predict the p-factor from stress-induced brain responses or FC changes.

#### Figures

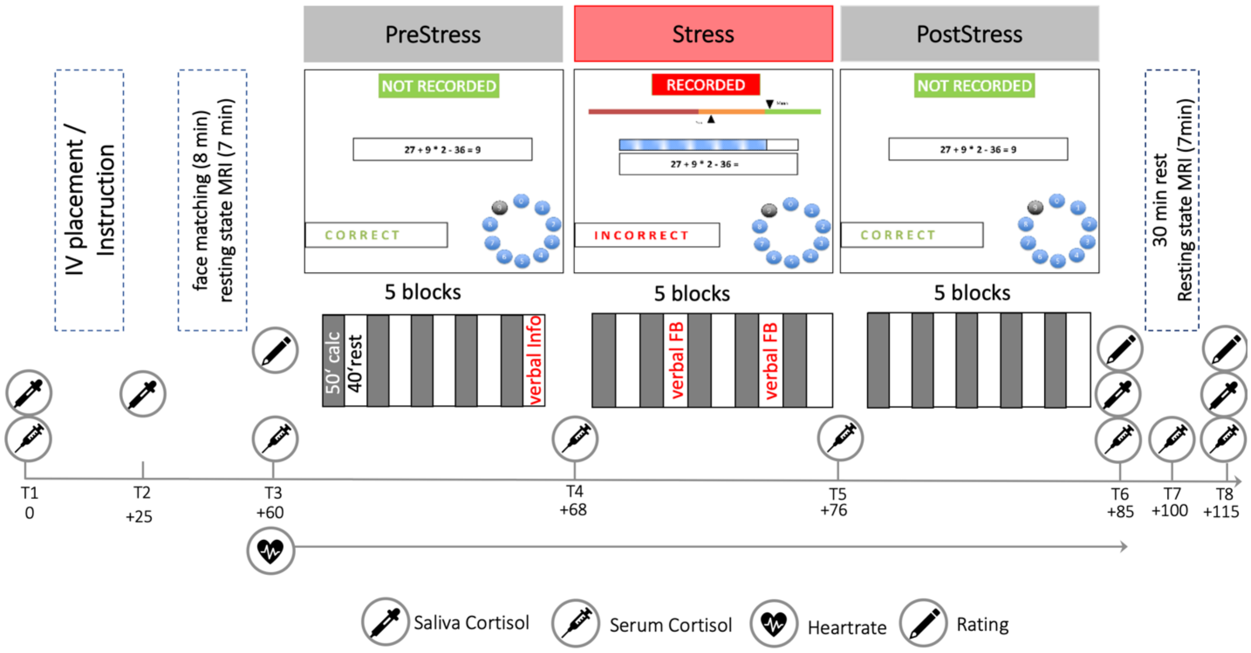


**Figure S1:** Detailed description of the psychosocial stress task (Kühnel et al., 2020). Before the stress phase, participants were informed about being recorded in the following trials. Additional aversive verbal feedback (verbal FB) about unsatisfactory performance was given in the 2nd and 4th rest period of the *Stress* condition. Saliva sampling was done in all participants (N=217) and in subsample of n=73 participants blood samples were taken to assess the cortisol response with higher temporal resolution.

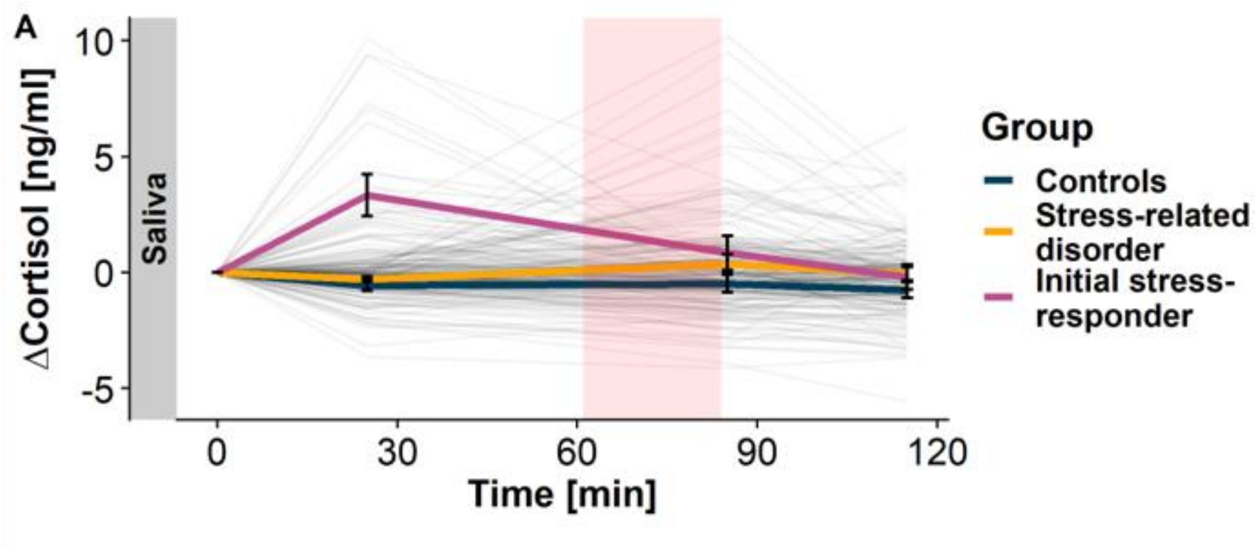


**Figure S2:** Trajectories of cortisol response differ between responders to the blood-drawing procedure in a subsample of 73 participants compared to participants without this initial response. The cortisol response was slightly albeit not significantly higher in participants with a mood- and/or anxiety disorder within the last 12-month ($b$=.24, $p$=.065) when excluding participants with a pre-task cortisol response.

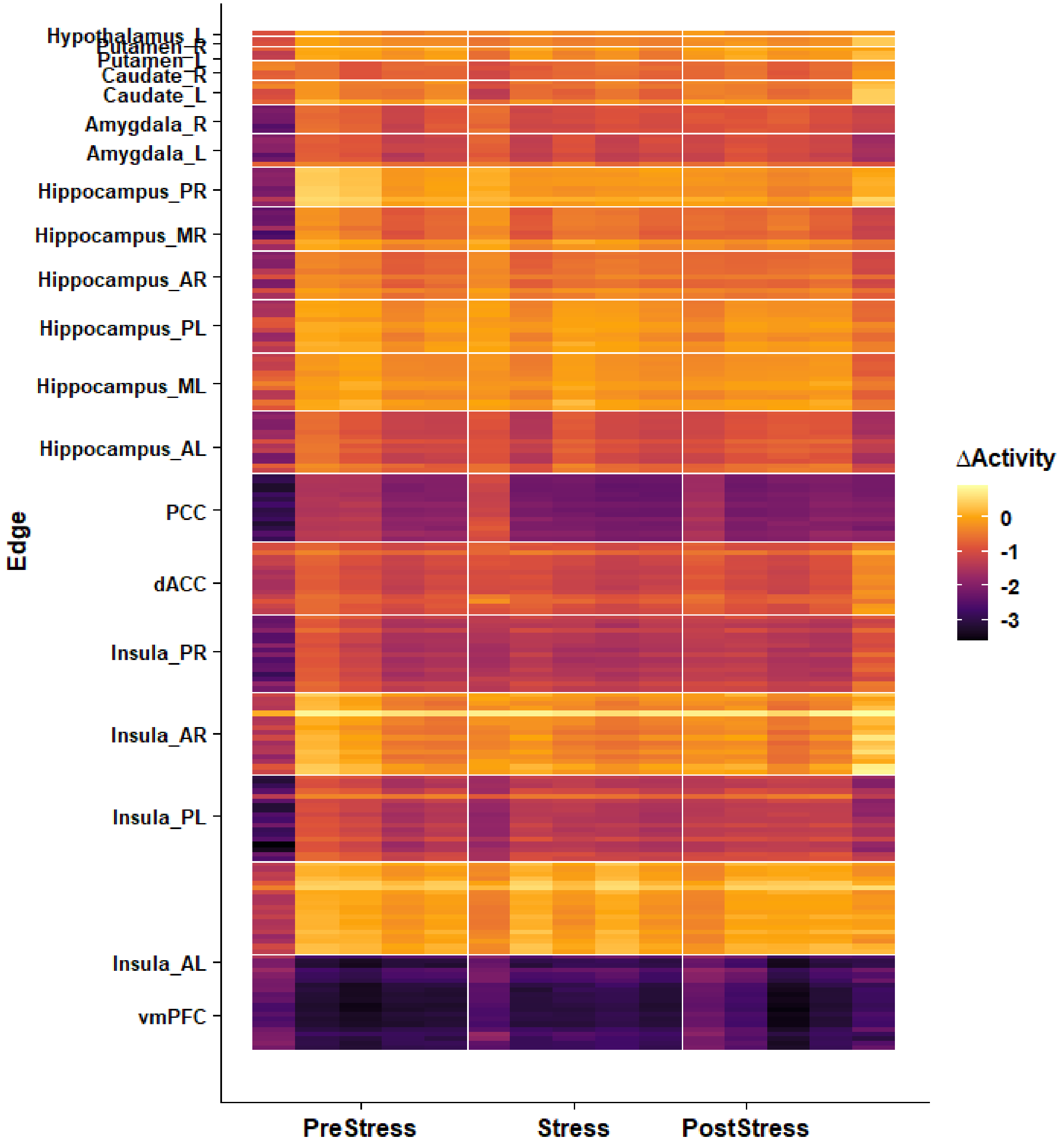


**Figure S3**. Changes in brain response (activation) across task blocks. Response magnitude is similar across the different phases of the stress cycle.

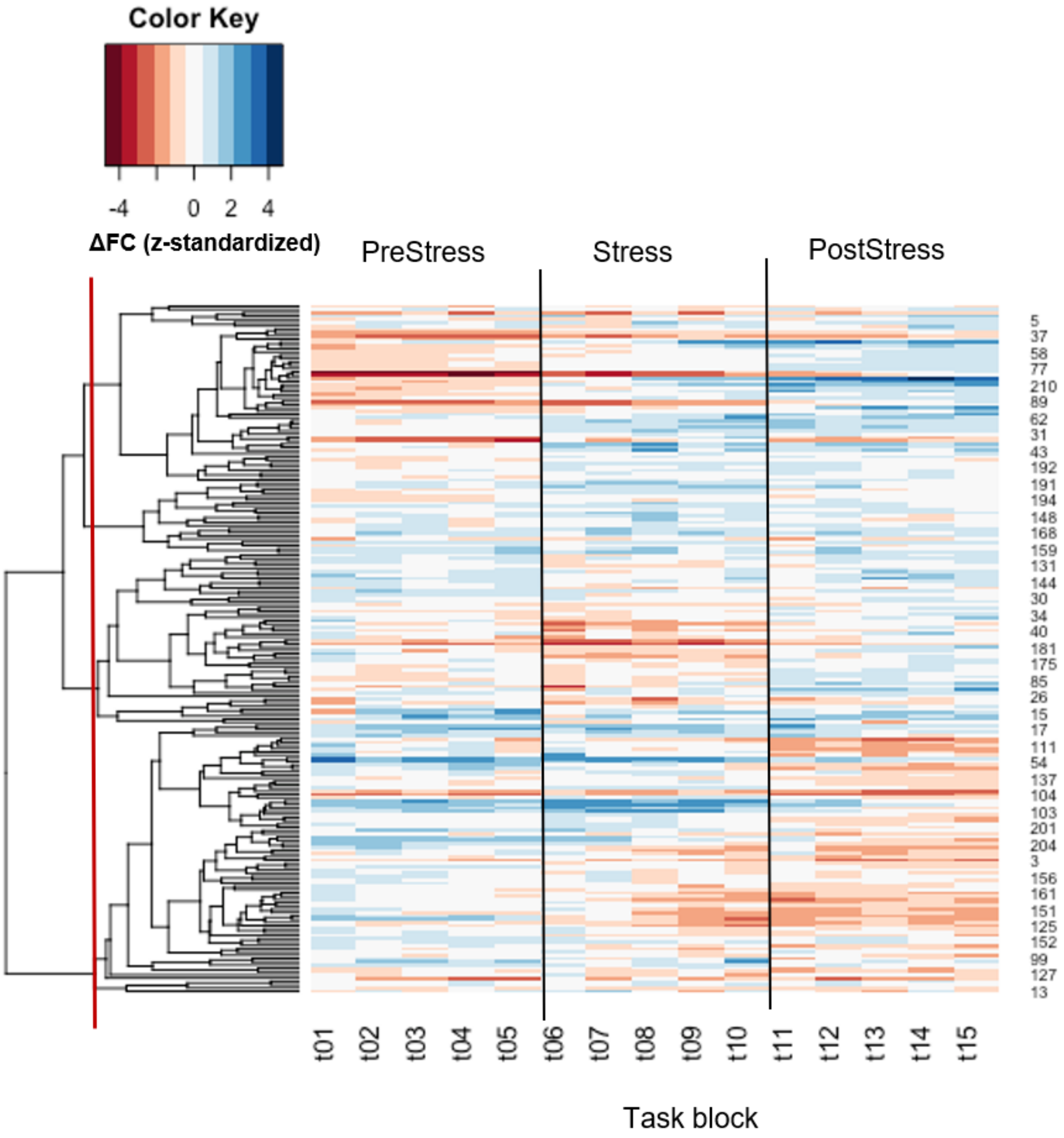


**Figure S4:** Dendrogram and ordered FC change matrix visualizing the hierarchical clustering solution. Choosing a more fine-grained resolution (i.e., more clusters) did not improve prediction of the intra- or inter-individual differences.

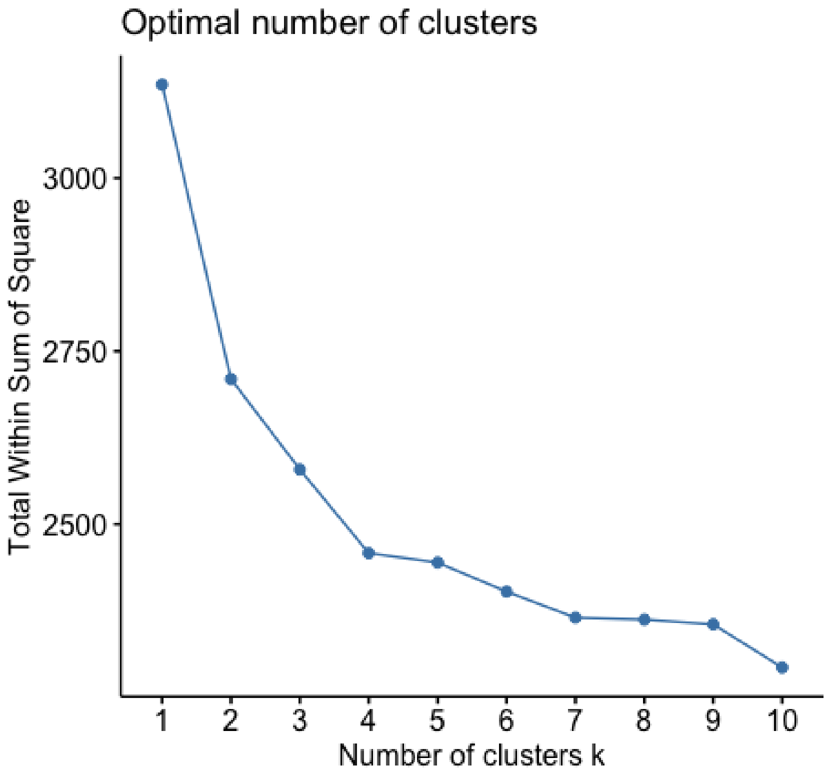


**Figure S5:** Scree plot for the total within cluster sum of squares (wss) for cluster solutions from k = 1-10. At 4 clusters the decrease in wss when adding additional clusters levels off.

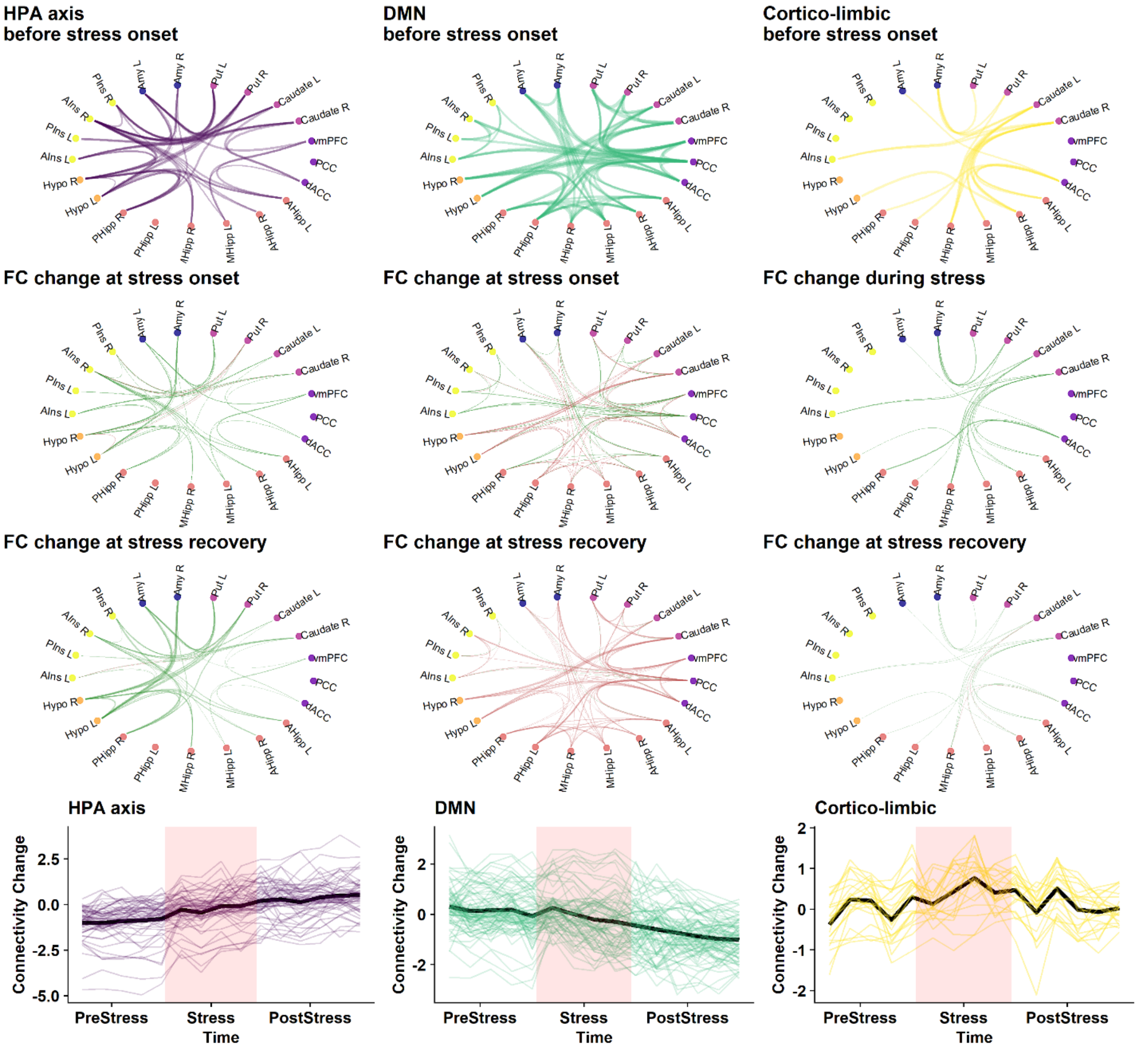


**Figure S6:** Circle plots for the three other subnetworks derived in the hierarchical clustering step with the corresponding average and single edge trajectories over time. The first row of circle plots (before stress onset) shows the functional connectivity (FC) change of each subnetwork compared to resting state FC. Line thickness corresponds to rescaled FC change. The second row shows stress-induced changes in FC compared to FC before stress onset (i.e., FC at stress onset (task-block 6) – FC at the beginning of the task (task-block 1). Line thickness corresponds to the rescaled magnitude of FC-change. Red lines show decreased FC and green lines increased FC. The third row shows the same difference but for the last task-block indicative of the recovery from stress. Animated circle plots showing change in FC strength with changing line thickness are available in the Supplementary material.

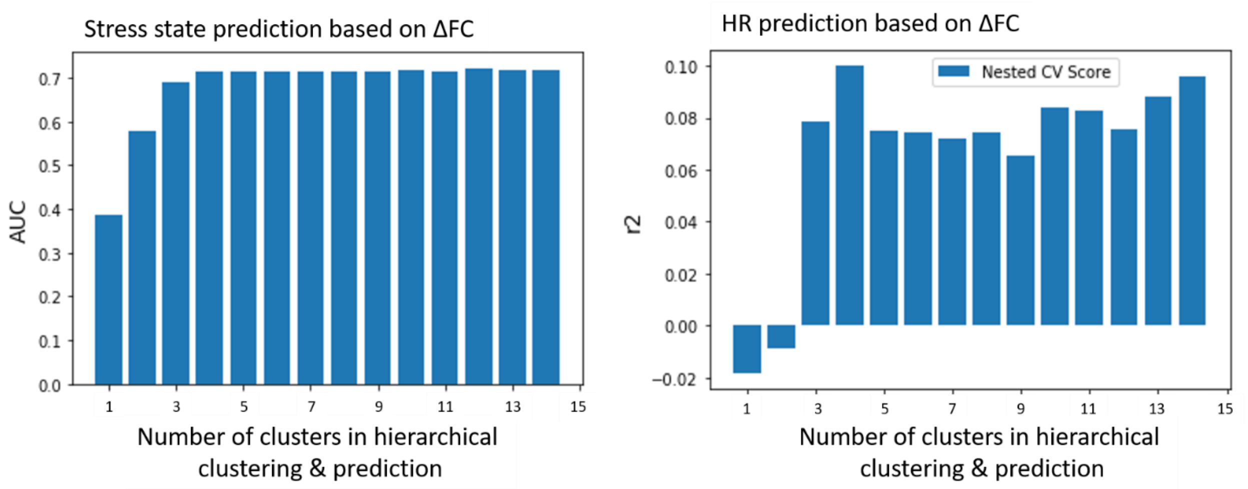


**Figure S7:** Predictive performance (nested cross-validation scores) for the intraindividual prediction of stress phase (AUC) and pulse rate change (R2) across different cluster solutions in the hierarchical clustering step. Performance does not improve when adding more than four clusters. Models were trained using support vector machine or support vector regressions.

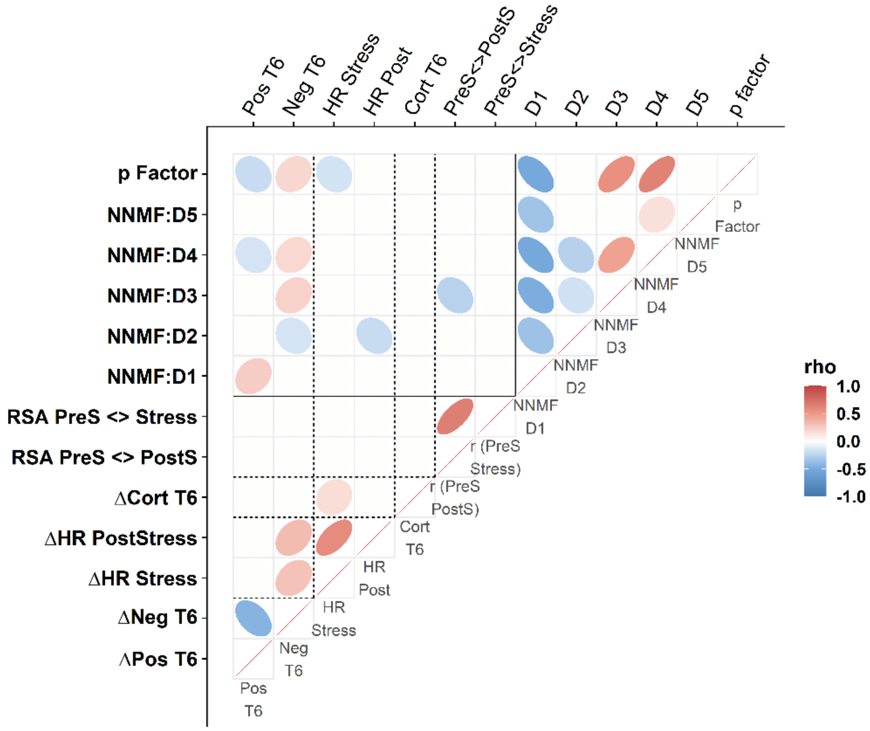


**Figure S8:** Correlation between the dimensions of symptoms of maladaptive stress reactivity derived using non-negative matrix factorization (NNMF) with autonomous, endocrine, and subjective stress reactivity to the psychosocial stress task. All correlations are partial correlations corrected for age, sex, and pre-task cortisol response (dummy coded yes/no). Only nominally significant correlations are shown (Supplementary Table S5). NNMF D1 = Resilience: Self-instruction, NNMF D2 = Resilience: Social support / cognitive, NNMF D3 = Intolerance of uncertainty, NNMF D4 = negative affectivity, NNMF D5 = Avoidance/Distraction, RSA PreS <> Stress = Neural similarity between *PreStress* and *Stress* condition, RSA PreS <> PostS = Neural similarity between *PreStress* and *PostStress* condition, ΔCort T6 = Cortisol increase after the end of the task (T6) compared to baseline (T0), ΔHR *PostStress* = Difference in pulse rate between task-block in the *PostStress* and *PreStress* condition, ΔHR Stress = Difference in pulse rate between task-block in the *PostStress* and *PreStress* condition, ΔNeg T6 = Difference in state negative affect directly after the task (T6) compared to before the task (T3), ΔPos T6 = Difference in state positive affect directly after the task (T6) compared to before the task (T3).

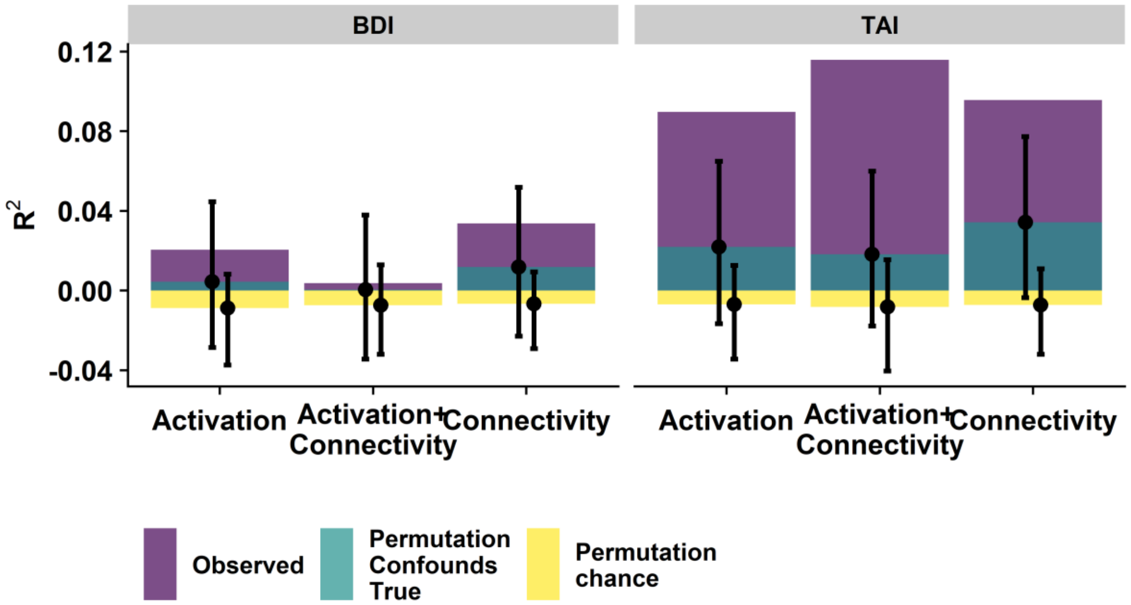


**Figure S9:** Trait anxiety (TAI) but not depressive symptoms (BDI) can be successfully predicted based on stress induced changes in functional connectivity and brain response and best when both are combined. Adding stress-induced changes in brain responses and FC improves the prediction of TAI compared to permutations of only the response variable (chance level, yellow) or the response variable and the confounding variables age, sex, average framewise displacement and pre-task cortisol response correspondingly (confound baseline, turquoise). Error bars depict 95% percentiles.

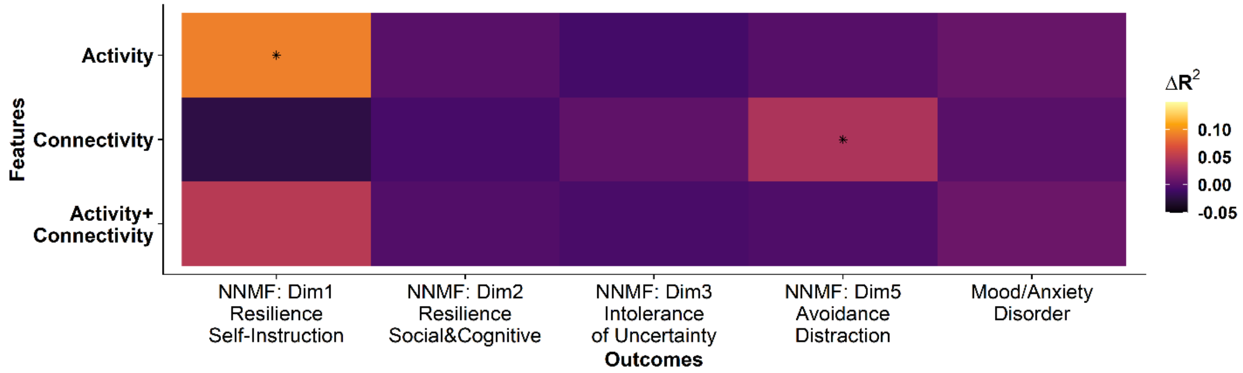


**Figure S10:** Predictive performance of the changes in FC or brain response for the other dimensions of psychological stress reactivity and the presence of any mood or anxiety disorder in the last 12 months. Adding stress-induced changes in brain responses but not FC improves the prediction of Resilience: Self-instruction compared to permutations of the response variable and the confounding variables age, sex, average framewise displacement and pre-task cortisol response correspondingly. Color indicates the increase in R2 compared to the prediction based on confounds. Asterisks indicate a one-sided p-value < .05.

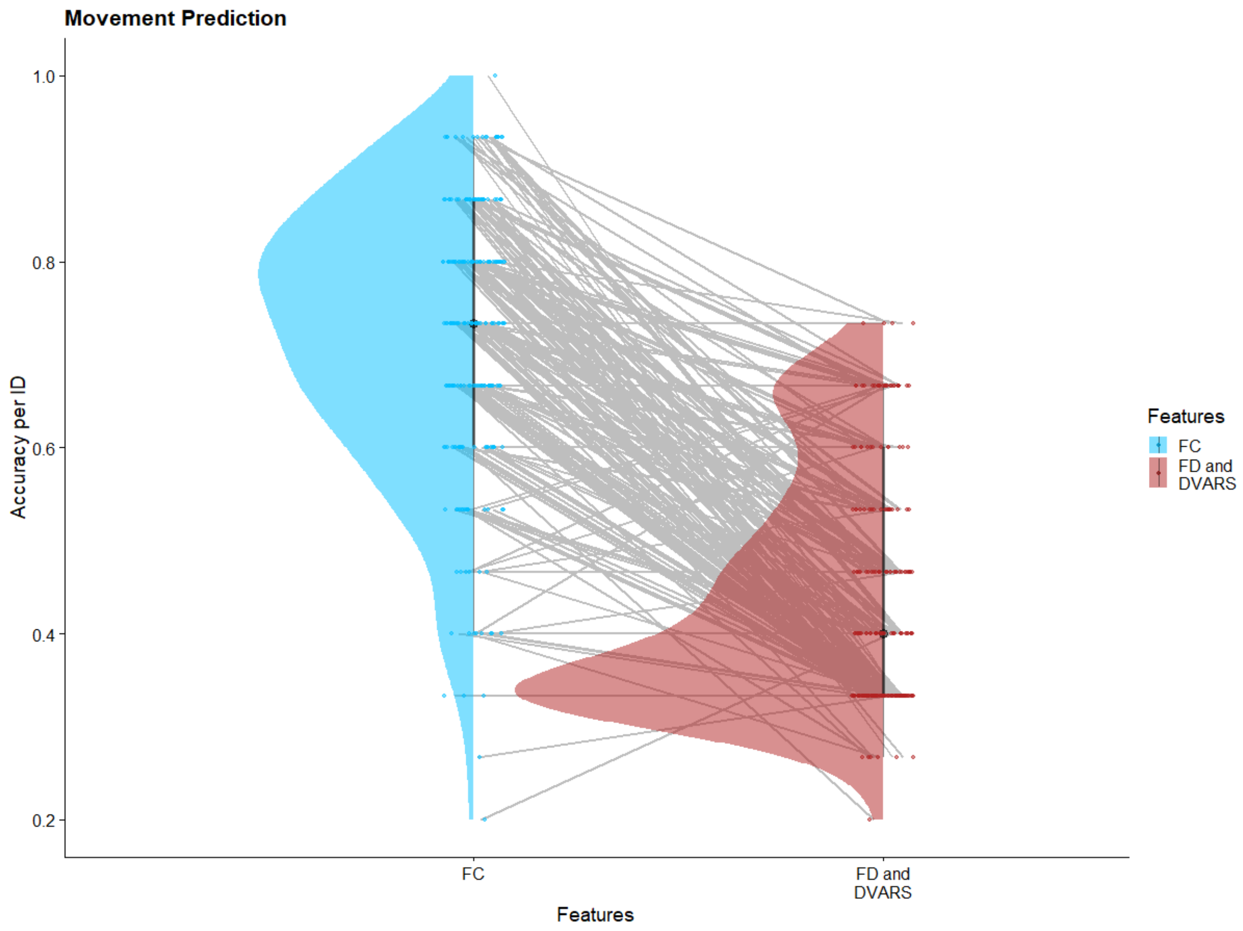


**Figure S11:** Prediction of the stress phase of a current task block is not driven by changes in head motion or image outliers. The accuracy of the prediction solely based on movement and DVARS estimates of each individual block only outperforms FC based prediction in 9 out of 221 individuals.

#### Tables

**Table S1**: Demographic and stress reactivity characteristics of sample and differences between participants with and without stress-related disorders within the last 12 months.

| Variable | N | Overall, N = 217[1] | 12-month healthy, N = 88[1] | Mood/Anxitey disorder, N = 129[1] | p-value[2] |
|---|---|---|---|---|---|
| **AGE** | 217 | 35.1 (12.1) | 33.3 (10.8) | 36.3 (12.8) | 0.13 |
| **SEX** | 217 | | | | 0.8 |
| Male | | 77 (35%) | 32 (36%) | 45 (35%) | |
| Female | | 140 (65%) | 56 (64%) | 84 (65%) | |
| **Pre-task cortisol** | 217 | | | | <0.001 |
| No | | 183 (84%) | 63 (72%) | 120 (93%) | |
| Yes | | 34 (16%) | 25 (28%) | 9 (7.0%) | |
| **BDI** | 196 | 12.5 (12.7) | 5.0 (7.5) | 17.8 (13.0) | <0.001 |
| **TAI** | 195 | 43.2 (14.8) | 33.1 (9.8) | 50.6 (13.6) | <0.001 |
| **SVF** | | | | | |
| **DIS** | 182 | 12.9 (4.2) | 13.6 (4.1) | 12.4 (4.2) | 0.11 |
| **SUBSA** | 182 | 9.9 (4.9) | 10.3 (5.0) | 9.7 (4.8) | 0.5 |
| **FLT** | 182 | 10.7 (6.4) | 8.6 (5.4) | 12.2 (6.6) | <0.001 |
| **RUM** | 182 | 14.9 (6.2) | 11.6 (5.8) | 17.3 (5.4) | <0.001 |
| **PLD** | 182 | 9.0 (5.4) | 12.0 (5.2) | 6.9 (4.5) | <0.001 |
| **POS** | 182 | 14.5 (5.4) | 16.1 (4.7) | 13.4 (5.6) | 0.001 |
| **REA** | 182 | 15.4 (4.2) | 15.3 (3.8) | 15.4 (4.5) | 0.9 |
| **RES** | 182 | 9.5 (5.9) | 6.9 (4.6) | 11.4 (6.0) | <0.001 |
| **GUI** | 182 | 10.4 (4.3) | 11.1 (4.3) | 9.9 (4.1) | 0.029 |
| **SEA** | 182 | 11.0 (5.8) | 8.6 (5.0) | 12.8 (5.7) | <0.001 |

| Variable | N | Overall, N = 217[1] | 12-month healthy, N = 88[1] | Mood/Anxitey disorder, N = 129[1] | p-value[2] |
|---|---|---|---|---|---|
| **SIT** | 182 | 15.5 (4.0) | 14.9 (3.8) | 16.0 (4.1) | 0.038 |
| **SOC** | 182 | 13.9 (5.5) | 14.1 (5.6) | 13.8 (5.5) | 0.6 |
| **AVO** | 182 | 12.9 (5.2) | 12.4 (5.2) | 13.3 (5.2) | 0.2 |
| **POS1** | 182 | 9.7 (4.0) | 11.6 (4.0) | 8.4 (3.5) | <0.001 |
| **POS2** | 182 | 11.4 (3.9) | 11.9 (3.9) | 11.0 (3.9) | 0.2 |
| **POS3** | 182 | 15.2 (3.8) | 15.4 (3.4) | 15.0 (4.1) | 0.4 |
| **NEG** | 182 | 11.6 (5.2) | 8.9 (4.3) | 13.4 (5.0) | <0.001 |
| **IOU** | 192 | 63.9 (23.7) | 50.7 (17.5) | 73.5 (23.0) | <0.001 |
| **Resilienz** | 192 | 60.6 (9.9) | 65.2 (7.8) | 57.3 (9.9) | <0.001 |

[1]Mean (SD); n (%), pre-task cortisol = cortisol response > .91 ng/ml at T1 (20 minutes after arrival) induced for example by the blood taking procedure in a subsample; BDI = Becks Depression inventory, TAI = trait anxiety inventory, SVF = stress coping questionnaire, DIS = distraction, SUBSA = substitutional satisfaction, FLT = flight tendency, RUM= rumination, PLD = playdown, POS = positive self-instruction, REA = reaction control, RES = resignation, GUI = guilt denial, SEA = self-accusation, SIT = situation control, SOC = need for social support, AVO = avoidance. POS1 – POS3 = stress-reducing strategies 1-3, NEG = stress-augmenting strategies, IOU = intolerance of uncertainty scale

[2]Wilcoxon rank sum test; Pearson's Chi-squared test

**Table S2**: Current and lifetime prevalence of psychiatric disorders identified using the CIDI in the present sample.

| | **12-months diagnosis N(%)** | **Lifetime diagnosis N(%)** |
|---|---|---|
| Substance use disorders (F1) | 10 (5%) | 47 (21%) |
| Mood disorders (F3) | 79 (36%) | 87 (40%) |
| Anxiety-related disorders (F4) | 119 (55%) | 142 (68%) |
| Other disorders | 10 (5%) | 18 (8%) |
| No diagnoses | 80 (37%) | 53 (24%) |
| 1 diagnosis | 67 (31%) | 65 (30%) |
| 2 diagnoses | 57 (26%) | 66 (31%) |
| 3 and more diagnoses | 12 (5%) | 32 (14%) |

*Note:* Anxiety disorders (F4) include specific phobias. Stress-related disorders in the last 12 months are defined as participants with a mood or anxiety-related disorder within the last 12 months excluding specific phobias. The control includes all other participants that might still receive diagnoses from other axes (e.g. substance use disorder: smoking).

**Table S3**: Stress reactivity on the endocrine, autonomous, neural, and subjective level does not differ in participants reporting at least one stress-related disorder within the last 12 months.

| Stress marker | N | β Mood/Anxiety Disorder (yes/no) | 95% CI[1] | p-value |
|---|---|---|---|---|
| ΔCortisol (T6 – T1) | 210 | 0.14 | -0.16, 0.43 | 0.36 |
| ΔNegative affect (T6) | 216 | 1.0 | -0.21, 2.2 | 0.10 |
| ΔPositive affect (T6) | 216 | -0.32 | -0.87, 0.23 | 0.25 |
| ΔNegative affect (T8) | 216 | 0.28 | -0.54, 1.1 | 0.50 |
| ΔPositive affect (T8) | 216 | -0.35 | -0.90, 0.20 | 0.21 |
| Pulse rate ΔStress [bpm] | 190 | -0.82 | -1.9, 0.23 | 0.12 |
| Pulse rate Δ*PostStress* [bpm] | 190 | -0.71 | -1.5, 0.09 | 0.082 |
| Similarity PreStress Stress | 217 | 0.00 | -0.04, 0.03 | 0.80 |
| Similarity PostS PreS | 217 | -0.05 | -0.09, -0.02 | 0.005 |

[1]CI = Confidence Interval. All coefficients are estimated using linear models that additionally include age, sex, and pre-task cortisol response as well as average log-transformed framewise displacement for neural similarity.

**Table S4**: Number of participants included across all analyses.

| ***N*** | Subjective | Endocrine | Autonomous | Neural | NNMF |
|---|---|---|---|---|---|
| Subjective | 216 | | | | |
| Endocrine | 209 | 210 | | | |
| Autonomous (Pulse rate) | 189 | 185 | 190 | | |
| Neural (fMRI) | 216 | 212 | 190 | 217 | |
| NNMF | 172 | 173 | 152 | 175 | 175 |

Note: Exclusion reasons: Endocrine (salivary cortisol) not enough material, Autonomous (Pulse rate) insufficient data quality; NNMF (Non-negative matrix factorization) missing questionnaire data (n=21 (BDI), n=22 (TAI), n=35 (Coping), n=25 (Resilience and intolerance of uncertainty)).

**Table S5**: Partial correlations (corrected for age, sex, and pre-task cortisol) between symptoms of maladaptive stress reactivity derived by non-negative matrix factorization (NNMF) and endocrine, autonomous, and subjective responses to the psychosocial stress task.

| | ∆Pos T6 | ∆Neg T6 | ∆HR Stress | ∆HR Post | ∆Cort T6 | RSA PostS <> PreS | RSA Stress <> PreS | NNMF D1 | NNMF D2 | NNMF D3 | NNMF D4 |
|---|---|---|---|---|---|---|---|---|---|---|---|
| ∆Pos T6 | . | x | x | x | x | x | x | x | x | x | x |
| ∆Neg T6 | -.44** | . | x | x | x | x | x | x | x | x | x |
| ∆HR Stress | -.13 | .30** | . | x | x | x | x | x | x | x | x |
| ∆HR Post | -.10 | .33** | .58** | . | x | x | x | x | x | x | x |
| ∆Cort T6 | -.07 | .08 | .16* | -.08 | . | x | x | x | x | x | x |
| RSA PostS <> PreS | .13 | -.08 | .01 | -.14 | -.02 | . | x | x | x | x | x |
| RSA Stress <> PreS | .07 | -.02 | .04 | -.09 | -.11 | .65** | . | x | x | x | x |
| NNMF D1 | .24** | -.11 | .04 | .00 | .01 | .10 | -.03 | . | x | x | x |
| NNMF D2 | .02 | -.16* | -.09 | -.20* | .01 | .01 | .02 | -.36** | . | x | x |
| NNMF D3 | -.13 | .21* | .04 | .15 | .00 | -.26** | -.14 | -.48** | -.17* | . | x |
| NNMF D4 | -.16* | .19* | -.13 | .00 | -.02 | -.13 | .04 | -.52** | -.26** | .46** | . |
| NNMF D5 | -.04 | .05 | .03 | .09 | -.09 | -.04 | .02 | -.35** | -.08 | .12 | .15 |

Note: * $p_{(uncorrected)} < .05$, ** $p_{(uncorrected)} < .001$, NNMF D1 = Resilience: Self-instruction, NNMF D2 = Resilience: Social support / cognitive, NNMF D3 = Intolerance of uncertainty, NNMF D4 = negative affectivity, NNMF D5 = Avoidance/Distraction, RSA PreS <> Stress = Neural similarity between *PreStress* and Stress condition, RSA PreS <> PostS = Neural similarity between *PreStress* and *PostStress* condition, ∆Cort T6 = Cortisol increase after the end of the task (T6) compared to baseline (T0), ∆HR *PostStress* = Difference in pulse rate between task-block in the *PostStress* and *PreStress* condition, ∆HR Stress = Difference in pulse rate between task-block in the *PostStress* and *PreStress* condition, ∆Neg T6 = Difference in state negative affect directly after the task (T6) compared to before the task (T3), ∆Pos T6 = Difference in state positive affect directly after the task (T6) compared to before the task (T3).

### References


Ashburner, J., 2007. A fast diffeomorphic image registration algorithm. NeuroImage 38, 95–113. https://doi.org/10.1016/j.neuroimage.2007.07.007

Behzadi, Y., Restom, K., Liau, J., Liu, T.T., 2007. A component based noise correction method (CompCor) for BOLD and perfusion based fMRI. NeuroImage 37, 90–101. https://doi.org/10.1016/j.neuroimage.2007.04.042

Caspi, A., Houts, R.M., Belsky, D.W., Goldman-Mellor, S.J., Harrington, H., Israel, S., Meier, M.H., Ramrakha, S., Shalev, I., Poulton, R., Moffitt, T.E., 2014. The p Factor: One General Psychopathology Factor in the Structure of Psychiatric Disorders? Clinical Psychological Science 2, 119–137. https://doi.org/10.1177/2167702613497473

Fried, E.I., Greene, A.L., Eaton, N.R., 2021. The p factor is the sum of its parts, for now. World Psychiatry 20, 69–70. https://doi.org/10.1002/wps.20814

Goldfarb, E.V., Seo, D., Sinha, R., 2019. Sex differences in neural stress responses and correlation with subjective stress and stress regulation. Neurobiology of Stress 11, 100177. https://doi.org/10.1016/j.ynstr.2019.100177

Janke, W., 1994. Befindlichkeitsskalierung durch Kategorien und Eigenschaftswörter: BSKE (EWL) nach Janke, Debus, Erdmann und Hüppe. Test und Handanweisung. Unveröffentlichter Institutsbericht, Lehrstuhl für Biologische und Klinische Psychologie der Universität Würzburg.

Janke, W., Debus, G., 1978. Die Eigenschaftswörterliste: EWL. Verlag für Psychologie CJ Hogrefe.

Kühnel, A., Kroemer, N.B., Elbau, I.G., Czisch, M., Sämann, P.G., Walter, M., Binder, E.B., 2020. Psychosocial stress reactivity habituates following acute physiological stress. Human Brain Mapping 41, 4010–4023. https://doi.org/10.1002/hbm.25106

Rodríguez-Liñares, L., Vila, X., Mendez, A., Lado, M., Olivieri, D., 2008. RHRV: An R-based software package for heart rate variability analysis of ECG recordings, in: 3rd Iberian Conference in Systems and Information Technologies (CISTI 2008). pp. 565–574.

Shen, X., Tokoglu, F., Papademetris, X., Constable, R.T., 2013. Groupwise whole-brain parcellation from resting-state fMRI data for network node identification. Neuroimage 0, 403–415. https://doi.org/10.1016/j.neuroimage.2013.05.081

Vest, A.N., Poian, G.D., Li, Q., Liu, C., Nemati, S., Shah, A.J., Clifford, G.D., 2018. An open source benchmarked toolbox for cardiovascular waveform and interval analysis. Physiol. Meas. 39, 105004. https://doi.org/10.1088/1361-6579/aae021

Vila, J., Palacios, F., Presedo, J., Fernandez-Delgado, M., Felix, P., Barro, S., 1997. Time-frequency analysis of heart-rate variability. IEEE Engineering in Medicine and Biology Magazine 16, 119–126. https://doi.org/10.1109/51.620503
